# Dissection of complex, fitness-related traits in multiple *Drosophila* mapping populations offers insight into the genetic control of stress resistance

**DOI:** 10.1101/383802

**Authors:** Elizabeth R. Everman, Casey L. McNeil, Jennifer L. Hackett, Clint L. Bain, Stuart J. Macdonald

**Author notes:** Corresponding author: Stuart Macdonald, Department of Molecular Biosciences, 4043 Haworth Hall, 1200 Sunnyside Avenue, University of Kansas, Lawrence, KS 66045.

## Abstract

We leverage two complementary *Drosophila melanogaster* mapping panels to genetically dissect starvation resistance, an important fitness trait. Using >1600 genotypes from the multiparental *Drosophila* Synthetic Population Resource (DSPR) we map numerous starvation stress QTL that collectively explain a substantial fraction of trait heritability. Mapped QTL effects allowed us to estimate DSPR founder phenotypes, predictions that were correlated with the actual phenotypes of these lines. We observe a modest phenotypic correlation between starvation resistance and triglyceride level, traits that have been linked in previous studies. However, overlap among QTL identified for each trait is low. Since we also show that DSPR strains with extreme starvation phenotypes differ in desiccation resistance and activity level, our data imply multiple physiological mechanisms contribute to starvation variability. We additionally exploited the *Drosophila* Genetic Reference Panel (DGRP) to identify sequence variants associated with starvation resistance. Consistent with prior work these sites rarely fall within QTL intervals mapped in the DSPR. We were offered a unique opportunity to directly compare association mapping results across labs since two other groups previously measured starvation resistance in the DGRP. We found strong phenotypic correlations among studies, but extremely low overlap in the sets of genomewide significant sites. Despite this, our analyses revealed that the most highly-associated variants from each study typically showed the same additive effect sign in independent studies, in contrast to otherwise equivalent sets of random variants. This consistency provides evidence for reproducible trait-associated sites in a widely-used mapping panel, and highlights the polygenic nature of starvation resistance.

## Introduction

Periods of food scarcity and suboptimal nutrient resources present an important challenge for nearly all species (McCue 2010), and this form of environmental stress can limit the survival of individuals with poor nutritional status and reduced stress resistance (Harshman *et al.* 1999; Lee and Jang 2014). As a result, starvation stress resistance has direct implications for the fitness of individuals as they experience resource variability in natural populations. Starvation resistance is a classic quantitative, fitness-related trait that is associated with several other phenotypes that influence survival, lifespan, and reproduction (Service and Rose 1985; Da Lage *et al.* 1990; Rose *et al.* 1992; Toda and Kimura 1997; Karan and Parkash 1998; Hoffmann *et al.* 2005b; Sørensen *et al.* 2007; Lee and Jang 2014). In particular, increased starvation resistance is often negatively correlated with fecundity and positively correlated with longevity, energy storage, and other stress response traits (Service *et al.* 1985; Rose *et al.* 1992; Hoffmann and Parsons 1993; Chippindale *et al.* 1993; Harshman *et al.* 1999; Bochdanovits and de Jong 2003; Bubliy and Loeschcke 2005; Sørensen *et al.* 2007; Schwasinger-Schmidt *et al.* 2012; Lee and Jang 2014). Because of pervasive phenotypic and genetic correlations between starvation resistance and these other traits, the evolution of starvation resistance in natural populations is complex, and is driven by adaptation to heterogeneous environments, phenotypic plasticity, and extensive pleiotropy (Service and Rose 1985; Hoffmann and Parsons 1991, 1993; Chippindale *et al.* 1993; Toda and Kimura 1997; Karan and Parkash 1998; Harshman *et al.* 1999; Harbison *et al.* 2004; Pijpe *et al.* 2007; Rion and Kawecki 2007; Bauerfeind *et al.* 2014; Colinet *et al.* 2015; Everman and Morgan 2018).

Artificial selection for starvation resistance often results in a concomitant increase in desiccation resistance (Hoffmann and Parsons 1989a; Chippindale *et al.* 1996; Hoffmann and Harshman 1999; Harshman *et al.* 1999; Hoffmann *et al.* 2001), and selection specifically on desiccation resistance can also result in a corresponding rapid increase in starvation resistance (Hoffmann and Parsons 1989b). Nonetheless, surveys of natural populations in several *Drosophila* species have shown that starvation and desiccation resistance can also independently vary (Davidson 1990; Karan and Parkash 1998; Karan *et al.* 1998; Chippindale *et al.* 1998; Hoffmann and Harshman 1999; Gilchrist *et al.* 2008; Goenaga *et al.* 2013). Given these variable patterns, Karan and Parkash (1998) and Da Lage *et al.* (1990) suggest that desiccation and starvation resistance may not routinely be associated. Rather, both traits may be directly and indirectly influenced by climate variability, and selection on other correlated traits such as diapause or thermal tolerance in seasonally variable temperate environments (Hoffmann and Parsons 1989b; Schmidt *et al.* 2005; Sørensen *et al.* 2007; Rion and Kawecki 2007; Goenaga *et al.* 2013; Rajpurohit *et al.* 2018).

Similar to desiccation, artificial selection for increased starvation resistance often results in an increase in lipid levels in *D. melanogaster* (Chippindale *et al.* 1996, 1998; Djawdan *et al.* 1998; Harshman *et al.* 1999; Schwasinger-Schmidt *et al.* 2012; Goenaga *et al.* 2013; Hardy *et al.* 2018), suggesting that energy storage is one important mechanism that contributes to starvation resistance. However, variation in the association between these traits has also been observed. For example, while Chippindale *et al.* (1996) provided evidence of a strong positive correlation between starvation resistance and lipid concentration following 60 generations of selection for starvation resistance, Hoffmann *et al.* (2001) found that total lipid concentration and starvation resistance in isofemale lines derived from natural populations were not correlated. Thus, the association between starvation resistance and lipid level is likely dependent upon genetic background and the evolutionary history of a population, resulting in across-population variation in the strength and direction of the correlation between these traits.

Genetic dissection of starvation resistance can both lead to the identification of loci impacting phenotypic variation and help understand how this trait is associated with desiccation resistance and lipid level. Several studies have examined the genetic basis of starvation resistance in *D. melanogaster* using a combination of selection experiments (Rose *et al.* 1992; Chippindale *et al.* 1996; Harshman *et al.* 1999; Bochdanovits and de Jong 2003; Bubliy and Loeschcke 2005; Schwasinger-Schmidt *et al.* 2012; Hardy *et al.* 2018; Michalak *et al.* 2018), gene expression studies following exposure to starvation stress (Harbison *et al.* 2005; Sørensen *et al.* 2007), and genetic mapping (Harbison *et al.* 2004; Mackay *et al.* 2012; Huang *et al.* 2014; Everman and Morgan 2018). These studies have provided extensive lists of candidate genes and variants, some of which have been functionally validated (Lin *et al.* 1998; Clancy *et al.* 2001; Harbison *et al.* 2004, 2005; Sørensen *et al.* 2007). However, up to this point few studies have undertaken an examination of the genetic architecture of triglyceride or lipid content in the same genetically diverse panel used to examine variation in starvation resistance. Doing so would allow a detailed comparison of quantitative trait loci (QTL) that contribute to variation in each trait, provide insight into the similarity of the genetic architectures of starvation resistance and correlated traits, and facilitate a better understanding of their evolution, and the mechanisms underlying their variation.

In this study we use two powerful *D. melanogaster* mapping panels - the *Drosophila* Synthetic Population Resource (DSPR) and the *Drosophila* Genetic Reference Panel (DGRP) - to genetically dissect phenotypic variation, and to explore the phenotypic and genetic relationships among traits, among mapping panels, and among laboratories. Our approach allows us to accomplish three primary objectives. First, by measuring starvation resistance and triglyceride level in the DSPR, we assess overlap in the loci that contribute to variation in each trait. Prior work on these traits in flies suggests they would show similar genetic architectures with many pleiotropic loci. However, despite a significant phenotypic correlation, we report limited overlap among mapped loci contributing to variation in starvation resistance and triglyceride level, suggesting that the genetic basis of these traits is largely independent in the DSPR. This highlights the role that other physiological mechanisms, such as activity level and desiccation resistance that also we explore here, may have in influencing starvation resistance.

Second, by measuring starvation resistance in both the DSPR and the DGRP under the same environmental conditions, we address variation in the genetic architecture of this trait between two distinct populations. In common with some previous studies using both panels to dissect a trait (e.g. Najarro *et al.* 2015, 2017), we also find little overlap in the loci associated with starvation resistance between mapping panels. This is likely the combined result of the populations having unique genetic backgrounds (King and Long 2017), distinct evolutionary histories, and differences in power to detect causative loci (Long *et al.* 2014).

Third, we leverage the ability to repeatedly measure trait variation on the same, stable set of inbred genotypes to compare our DGRP starvation data to two previous starvation resistance datasets collected by different laboratories (Mackay *et al.* 2012; Everman and Morgan 2018). We found that the sign of the additive effects of the most strongly-associated SNPs were consistent across datasets. This suggests these SNPs contribute to variation in starvation resistance in the DGRP, but have sufficiently small effects that they are regularly not identified following genomewide multiple testing correction. This across-study comparison of the genetic architecture of starvation resistance provides both technical insight into the use of genomewide association (GWA) studies to understand the genetic basis of complex traits, and biological insight into the phenotypic effects of loci that contribute to trait variation.

## Materials and Methods

### Mapping populations

#### Drosophila *Synthetic Population Resource*

The DSPR is a multiparental population that consists of two synthetic populations (pA and pB) that were each established following an intercross of eight highly-inbred founder lines, with one founder line shared between the two populations (King *et al.* 2012a). Flies were maintained in pairs of subpopulations (pA1, pA2, pB1, pB2) at high population density for 50 generations prior to the establishment of >1600 genotyped recombinant inbred lines (RILs) via 25 generations of full-sib inbreeding (King *et al.* 2012a; b). Founder lines for the pA and pB panels have also been sequenced at 50x coverage, enabling inference of the haplotype structure of each RIL via a hidden Markov model (described in King *et al.* (2012a)).

#### Drosophila *Genetic Reference Panel*

The DGRP was established from mated females collected from a natural population in Raleigh, North Carolina, with inbred lines derived from 20 generations of full-sib mating (Mackay *et al.* 2012). Each of the 205 DGRP lines have been re-sequenced and genotyped allowing GWA mapping to be carried out in the panel (Mackay *et al.* 2012; Huang *et al.* 2014).

### Phenotyping assays and analysis

#### Large-scale starvation resistance assay

Strains from the DSPR and DGRP were duplicated from stocks, and flies were allowed to lay eggs for up to 2 days. Vials were inspected twice daily, and laying adults were cleared when necessary to maintain a relatively even egg density across experimental vials. While this visual method of density control is less precise than counting eggs, experiments with 20 randomly-selected DSPR RILs showed that the effect on starvation resistance of rearing flies via egg counting or by visually-assessing egg number is extremely limited (variance explained = 0.9%; Fig S1; see Table S1 for full breakdown of variance components).

In the following generation, experimental flies (2-4 days old) were sorted by sex over light CO_2_ anesthesia and placed in groups of same-sex individuals on new cornmeal-molasses-yeast media for 1 day until the start of the starvation assay. The assay was initiated by placing flies on 1.5% agar media that additionally contained preservatives - a mix of propionic and phosphoric acids, and benzoic acid (Tegosept, Genesee Scientific) dissolved in ethanol (see starvation media recipe Text S1). Starvation media was made within 24 hours of the initiation of each block of the assay and was not replaced throughout its duration. Vials were barcoded during the screen, blinding experimenters to strain identification number, and assisting with efficient data collection and analysis.

Death was assessed for each vial twice per day at approximately 0900 and 2100 hrs. The first assessment of survival was made 24 hours after flies were transferred to starvation media. Dead flies at this initial assessment point were not included in the analysis as their death may have resulted from handling during the initial transfer to experimental vials rather than starvation stress. Vials containing flies that had become entangled in the cotton vial plug at any point during the assay were also excluded from the analysis. Flies were considered dead if they were not moving or were unable to dislodge themselves from the starvation media. The phenotype used for mapping was the mean time to death in hours per strain across the vial replicates. Flies for this screen were reared and tested at approximately 23°C, 30-60% humidity, with constant light.

We screened the DSPR (861 pA1/pA2 and 864 pB1/pB2 RILs) in a series of batches across a seven-month period in 2010. Each batch included the majority of RILs that belonged to a particular subpopulation. Starvation resistance was measured in 168 DGRP lines in a single batch in 2012. In both mapping panels, survival was measured across 2 vial replicates per sex in ~85% of strains, with ~90% of vials containing 10 flies (minimum flies per vial = 6). Finally, we measured starvation resistance in the 15 DSPR founder lines, using 5 vial replicates per founder, in one batch.

We assessed variation in starvation resistance due to subpopulation and sex in the DSPR with a two-way ANOVA, including the interaction, and treated subpopulation (pA1, pA2, pB1, pB2) and sex as fixed factors. Male and female-specific differences among the four subpopulations were tested using Tukey’s HSD post hoc comparisons with an experiment-wide α = 0.05. Differences in starvation resistance due to sex among the DGRP lines were analyzed with a one-way ANOVA, treating sex as a fixed factor.

#### Desiccation resistance assay

To investigate the correlation between starvation and desiccation resistance, we measured desiccation resistance in a subset of pA1/pA2 RILs that exhibited very low (17 RILs) or very high (16 RILs) average female starvation resistance in the large-scale screen. Desiccation resistance of female flies from all 33 strains was assessed in a single batch with two vial replicates per RIL, where 92.9% of vials contained 10 flies (minimum flies per vial = 8). We placed experimental flies, reared as described above, in empty vials plugged with cotton inside an airtight desiccator (Cleatech, LLC). Relative humidity was reduced to < 5% throughout the experiment by adding a large quantity of Drierite (calcium sulfate) to the chamber. Survival was assessed every hour following initiation of the experiment, and mean desiccation resistance per RIL was used in all analyses.

#### Activity assay

We employed the *Drosophila* Activity Monitoring System (DAM2, TriKinetics, Inc.) to assess activity levels both prior to, and during starvation for a subset of DSPR RILs, selecting 16 (19) pA1/pA2 RILs that exhibited high (low) female starvation resistance in the large-scale screen. Sixteen flies of each sex were tested per RIL. Flies for these assays were reared and tested at 25°C, 50% relative humidity, with a 12:12hr light:dark photoperiod. These environment conditions are different from our large-scale screen, but in line with those used in previous starvation resistance studies in *D. melanogaster* (Mackay *et al.* 2012; Everman and Morgan 2018). This change allowed us to examine the stability of DSPR starvation phenotypes across assay environments.

One day prior to adding flies to monitor tubes, cornmeal-yeast-dextrose media was poured into 100mm diameter petri dishes and allowed to set. Polycarbonate activity monitor tubes (5mm diameter × 65mm length) were filled to approximately 10mm by pushing them into the media, and the food plug in each tube was sealed with paraffin wax. A single fly was aspirated into each tube, and the tubes were capped with small pieces of Droso-Plugs (Genesee Scientific). Flies were allowed to acclimate to the tubes for 24 hours, and then we measured activity for the next 24 hours under non-stressful conditions. Subsequently, each fly was tipped to a second monitor tube containing starvation media (Text S1) and activity was continuously monitored until each fly died.

To determine differences in activity under non-stressful conditions due to starvation resistance rank (high versus low), we used a full three-way ANOVA model with interactions, and treated starvation rank, sex, and light status (light versus dark) as fixed effects. The effect size of the main effects and interactions were calculated using Cohen’s F, which determines the effect size as a ratio of the between-group and between-replicate standard deviations (R package: sjstats) (Cohen 1988; Quinn and Keough 2002; Lüdecke 2018).

#### Triglyceride level assay

We duplicated 311 pA1/pA2 and 628 pB1/pB2 DSPR RILs from stocks to two replicate vials, clearing parental flies when necessary to maintain relatively even egg density over test vials. Eleven days following the start of egg laying we collected two sets of 10-12 females from each parental vial, resulting in four collection vials from each RIL. Flies were aged for three additional days before measuring triglyceride level.

Experimental females from each collection vial were anesthetized using CO_2_, and groups of 5 were arrayed into deep well plates (Axygen, P-DW-11-C) over ice, with each well pre-loaded with a single glass bead. This resulted in 8 replicate samples of 5 females per RIL. Immediately after finishing a plate, we added 400μl of cold homogenization buffer (10mM potassium phosphate monobasic, 1mM EDTA, 0.05% Tween-20) to each well, homogenized for 45sec in a Mini-BeadBeater-96 (Bio Spec Products, Inc.), and centrifuged for 4min at 2,500g. We then moved 50μl of the supernatant to a standard PCR plate, incubated the plate in a thermocycler at 70°C for 5min, and then placed the plate on ice for 5min.

During the incubation steps, we added 30μl of homogenization buffer to 92 of the 96 wells of a flat-bottom, polystyrene assay plate (Greiner, 655101), and subsequently added 20μl of the heat-deactivated fly homogenate to each. The four remaining wells of every assay plate were dedicated to controls; one blank well contained 50μl of homogenization buffer only, and three wells contained 5μl of Glycerol Standard Solution (SigmaAldrich, G7793, 2.5mg/ml) along with 45μl of homogenization buffer.

The assay plate was then inserted into a BioTek Powerwave XS2 instrument pre-heated to 37°C and read at 540nm (baseline absorbance scan). After the scan, and within 10min, we added 100μl of Free Glycerol Reagent (SigmaAldrich, F6428) to each well. The plate was then re-inserted into the instrument, incubated at 37°C for 5min, and read again at 540nm (free glycerol absorbance scan). After this second scan, and again within 10min, we added 25μl of Triglyceride Reagent (SigmaAldrich, T2449) to each well. The plate was again incubated at 37°C for 5min in the machine and read for a third time at 540nm (triglyceride, or final absorbance scan).

For each sample, we obtained the final absorbance for each sample (FA_sample_) and calculated the initial absorbance (IA_sample_) as the free glycerol measurement minus the baseline measurement. We also generated the average final absorbance for the three standard wells (FA_std_) and the initial absorbance for the one blank well (IA_blank_). We then estimated the true serum triglyceride level as

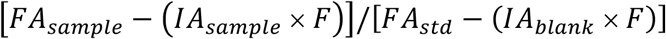

where *F* = 0.8. We then multiplied this value by the concentration of the glycerol standard solution (2.5mg/ml) and used the average value across all 8 replicate samples as the RIL mean triglyceride level for mapping and analysis. For precise details of the enzyme assay and triglyceride calculation, see the SigmaAldrich Serum Triglyceride Determination kit product insert (TR0100). Differences in triglyceride level due to DSPR subpopulation were investigated with a one-way ANOVA followed by *post hoc* comparisons using Tukey’s HSD (experiment-wide α = 0.05).

#### Correlations among traits

We assessed the relationship between the DSPR RIL mean starvation phenotypes from the large-scale screen with those from the activity monitor experiment, the desiccation resistance measures, and triglyceride level using general linear models. Subpopulation (pA1, pA2, pB1, pB2) was included as a factor in the analysis examining starvation resistance and triglyceride content.

Correlations among three DGRP starvation datasets - that from Mackay et al. (2012), Everman and Morgan (2018), and the new screen we report here - were examined in a pairwise manner using line means, accounting for multiple comparisons with a Bonferroni-adjusted alpha level. Differences in the overall mean starvation resistance among the three datasets were analyzed with a one-way ANOVA, treating study as a fixed factor.

#### Heritability

The genetic and phenotypic variances of starvation resistance and triglyceride content for the pA and pB DSPR panels, and of starvation resistance for the DGRP panel, were estimated with a linear mixed model using the lme and varcomp functions in R (R package: APE, Paradis *et al.* 2004; R package: nlme, Pinheiro *et al.* 2017). We calculated broad-sense heritability as the proportion of the total variance of the strain-specific response explained by the estimated genetic variance component (Lynch and Walsh 1998).

#### QTL mapping in the DSPR

Methods for QTL analysis, and the power and resolution of mapping using the DSPR panel are discussed in detail in King *et al.* (2012a,b). Briefly, QTL mapping and peak analysis were executed for starvation resistance and triglyceride data using the DSPRqtl R package (github.com/egking/DSPRqtl; FlyRILs.org), regressing the mean trait response for each RIL on the additive probabilities that each of the 8 founders contributed the haplotype of the RIL at each mapped position. Significance thresholds were assigned following 1000 permutations of the data, and positions of putative causative loci were estimated with 2-LOD support intervals, which approximate 95% confidence intervals for QTL position in the DSPR (King *et al.* 2012a). Mean starvation resistance varied between sexes in both the pA and pB panel (F_3,3440_ = 18.317; p < 0.0001; Fig S2; Table S2), and subpopulation influenced female starvation resistance in the pA panel (Tukey’s HSD p < 0.0001; Fig S2). Therefore, QTL mapping was performed for males and females of each panel separately, and subpopulation was included as a covariate in the analysis of pA females. Mean female triglyceride level was similar between the pA1 and pA2 subpopulations (Tukey’s HSD p = 0.75; Fig S3; Table S3) but varied between the pB1 and pB2 subpopulations (Tukey’s HSD p < 0.0001; Fig S3; Table S3), so subpopulation was included as a covariate in QTL analysis of the pB triglyceride data.

#### Analysis of DGRP starvation data

Variants associated with male and female starvation resistance in the DGRP were identified using the DGRP2 web-based GWA mapping tool (http://dgrp2.gnets.ncsu.edu), which takes into account variable *Wolbachia* infection status and large inversions that segregate among the lines (Mackay *et al.* 2012; Huang *et al.* 2014). We performed GWA analysis on data collected in this study and additionally reanalyzed starvation data from Mackay *et al.* (2012) and starvation of young flies (5 – 7 days old) from Everman and Morgan (2018). We additionally assigned the 150 DGRP lines that are shared between the three datasets an across-study mean and performed GWA analysis on this summary measure of starvation resistance.

SNPs associated with starvation resistance were identified within each of the four datasets following FDR correction for multiple comparisons (Benjamini and Hochberg 1995) in R (p.adjust; R Core Team 2017). Since we found no significantly associated SNPs with an FDR adjusted p-value < 0.05 for any starvation resistance dataset in either sex, we relaxed the significance threshold to an FDR adjusted p-value < 0.2. As a significance threshold of P < 10^−5^ is commonly used in the DGRP (e.g. Mackay *et al.* 2012; Morozova *et al.* 2014; Huang *et al.* 2014; Everman and Morgan 2018), we also present variants associated with starvation resistance in each of the four datasets using this threshold.

There was minimal overlap in the identity of the above-threshold, starvation-associated variants in each study. Thus, we sought to examine whether the sign of the additive effects of these sets of variants was preserved across studies. Additive effects were calculated as one-half the difference in starvation resistance between lines homozygous for the major allele and lines homozygous for the minor allele (major allele frequency > 0.5), after accounting for *Wolbachia* infection and TE insertions (Falconer and Mackay 1996; Huang *et al.* 2014). To determine the proportion of SNPs that are expected by chance to have additive effects of the same sign across studies, we obtained random samples of 50 SNPs from all of the DGRP SNP calls (~ 2 million SNPs) and calculated the additive effects of the sampled SNPs across pairs of datasets for each sex. To account for the possibility that the frequency spectrum of above-threshold (P < 10^−5^), associated SNPs is not represented by a set of randomly-selected variants, we stratified the random subsets of 50 SNPs according to the distribution of allele frequencies of the top 50 SNPs associated with starvation resistance for each sex in each study. Allele frequency bins used in this stratification were 0.05 – 0.1, > 0.1 – 0.2, > 0.2 – 0.3, > 0.3 – 0.4, and > 0.4 – 0. 5. The exact stratification for each sex and dataset is provided in Table S4. This process was repeated 1000 times for each paired comparison of datasets (6 comparisons total) using an ordinary nonparametric bootstrapping procedure with the R package boot (Davison and Hinkley 1997; Canty and Ripley 2017). For each iteration, we used a custom R function (see File S1) to calculate the proportion of the 50 random stratified SNPs that had positive additive effects in both of the datasets being compared.

#### Data availability

DGRP data from Mackay *et al.* (2012) are available online from http://dgrp2.gnets.ncsu.edu, and DGRP data from Everman and Morgan (2018) are available from Dryad (DOI: https://doi.org/10.5061/dryad.vq087). Data collected in this study is available from Dryad (available upon acceptance), including all raw data for starvation resistance in the DSPR and DGRP, raw desiccation resistance, triglyceride level, and activity data collected using the DSPR, and all mapping results (see File S2). R code for bootstrapping analysis and additive effect calculations in the DGRP is available in File S1.

## Results and Discussion

### Extensive phenotypic variation in starvation resistance in the DSPR and DGRP

Starvation resistance in both the DSPR and DGRP was highly variable among strains (Fig 1, Fig 2), and the broad sense heritability for starvation resistance was routinely high (Table 1). Males were typically less starvation resistant than females (Fig S2, Fig S4, Table S2, Table S5), although despite this male and female starvation resistance were significantly correlated in both the DSPR (pA: R^2 =^ 53.0%; pB: R^2 =^ 57.0%; Fig S5) and DGRP (R^2 =^ 68.0%; Fig S6). Such sex-specific differences in starvation resistance are likely influenced by a combination of higher glycogen and triglyceride levels and larger body size, which are often observed for females relative to males (Chippindale *et al.* 1996; Toda and Kimura 1997; Schwasinger-Schmidt *et al.* 2012; Goenaga *et al.* 2013).

**Table 1.** Broad sense heritability for starvation resistance and triglyceride level.

| Panel | Trait | Sex | Heritability |
| --- | --- | --- | --- |
| DSPR pA | Starvation | Female | 80.0% |
| DSPR pA | Starvation | Male | 73.1% |
| DSPR pB | Starvation | Female | 74.4% |
| DSPR pB | Starvation | Male | 71.5% |
| DGRP | Starvation | Female | 87.1% |
| DGRP | Starvation | Male | 87.0% |
| DSPR pA | Triglycerides | Female | 77.5% |
| DSPR pB | Triglycerides | Female | 82.3% |

**Figure 1.**
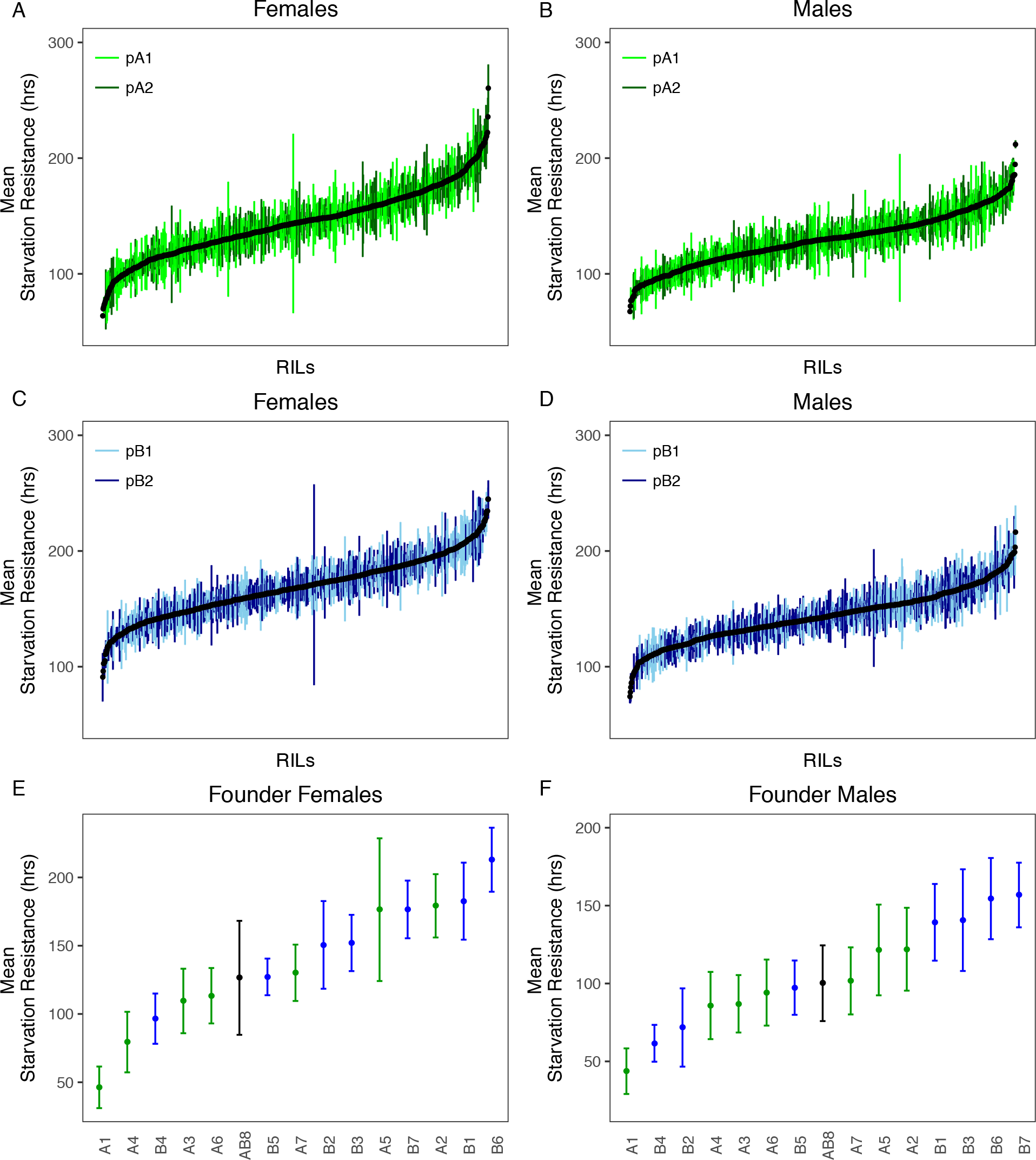
Variation in mean ((± SD) starvation resistance for each sex. A-D shows data for DSPR RIL panels pA and pB. E and F show data for the founder lines. In E and F, names of the founder lines are shown on the x-axis; the founder line AB8 is the founder shared by the two mapping panels.

**Figure 2.**
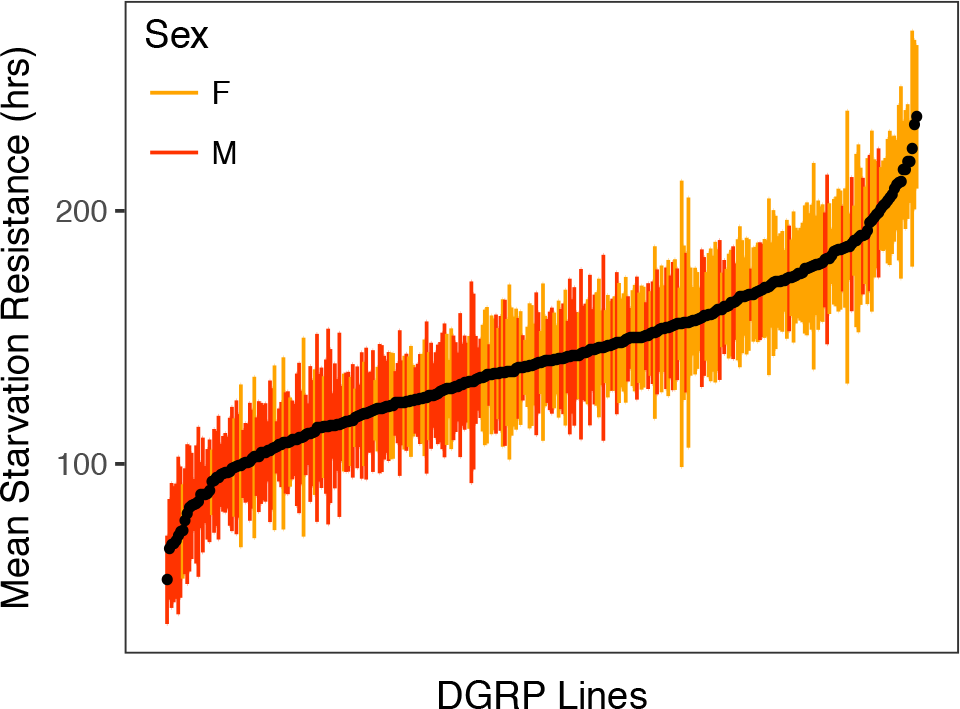
Variation in mean DGRP (±SD) starvation resistance in females (orange) and males (red).

We screened the DSPR and DGRP for starvation resistance at 23°C and under constant light conditions. Since starvation resistance is sensitive to the thermal environment (van Herrewege and David 1997; Karan and Parkash 1998; Karan *et al.* 1998; Hoffmann *et al.* 2005a; Bauerfeind *et al.* 2014) and may vary under different photoperiods (Sheeba *et al.* 2000; Xu *et al.* 2008; Seay and Thummel 2011), we sought to re-measure starvation resistance for a subset of DSPR RILs at 25°C and with a 12 hour: 12 hour light/dark cycle, conditions that have been used in other starvation studies (e.g. Mackay *et al.* 2012; Everman and Morgan 2018). Overall, starvation resistance of the re-tested RILs was lower in both sexes compared to that measured in the original large-scale starvation assay (effect of assay: F_1,136_ = 31.60, p < 0.0001; Fig S7A). Despite this, starvation resistance in the subset of RILs was significantly correlated between the two experiments (Females: β = 0.43±0.04, t = 9.7, p < 0.0001, R^2 =^ 73.9%; Males: β = 0.59±0.05, t = 10.9, p < 0.0001, R^2 =^ 78.3%; Fig S7B).

We similarly compared starvation resistance phenotypes for the DGRP measured in the current study with data generated by Mackay *et al.* (2012) and Everman and Morgan (2018). In our study, the DGRP exhibited considerably higher resistance than in these previous works (F_2,532_ = 1457.5, p < 0.0001; Fig S8). This discrepancy was not due to differences across studies in the frequency with which flies were counted (every 4, 8, or 12 hours depending on the study, Fig S8). To investigate whether the difference was due to the environmental conditions experienced by the experimental animals, we raised and tested 12 randomly-selected DGRP lines under the same conditions as described for our initial screen (i.e., 23°C, 30-60% relative humidity, and constant light) and under conditions that more closely mimic those described in Mackay *et al.* (2012) and Everman and Morgan (2018) (i.e. 25°C, 50% relative humidity, and 12:12hr light:dark). Furthermore, for both environments, we assayed starvation on agar media containing preservatives (see Text S1), and on media lacking preservatives, as used by Everman and Morgan (2018) and Mackay *et al.* (2012). The inclusion of preservatives in the assay media had the largest effect on variation in starvation resistance among studies (Preservatives: F_1,327_ = 1628.9, p ≪ 0.0001; variance explained = 81.2%; Fig S9), with rearing/testing environment explaining very little of the variation (see Table S6 for the full breakdown of ANOVA variance components). We speculate that the antibiotic properties of the preservatives extend lifespan under starvation conditions by limiting growth of pathogenic microorganisms.

Even given the large across-study difference in mean starvation resistance in the DGRP, we found moderately strong correlations in both sexes over datasets, ranging from 50.8% to 64.4% (Fig S10). The high correspondence among these three DGRP datasets, coupled with the phenotypic correlation between the subset of DSPR strains assayed using two different approaches (see above), suggests that fundamental aspects of the genetic control of starvation resistance are generally consistent over experiments, even when environmental conditions such as temperature are quite different. The differences we observe in starvation resistance between studies may reflect ecologically-relevant phenotypic plasticity. The temporally variable thermal environment is a particularly important source of selection for ectothermic organisms (Bell 2010; Bergland *et al.* 2014). Plastic shifts in starvation resistance in response to temperature can have important fitness benefits, including seasonal adaptation to fluctuating resource availability as has been reported in the butterfly *Bicyclus anynana* (Pijpe *et al.* 2007) and following the induction of diapause in *D. melanogaster* (Schmidt *et al.* 2005; Rion and Kawecki 2007). Collectively, these previous studies and our data speak to the important influence of both phenotypic plasticity and genotype on variation in starvation resistance in natural populations.

### Starvation resistance is associated with desiccation resistance and low activity in the DSPR

Environmental stress can exert selection pressure on energy use and storage, and environmental stressors that impact one type of stress resistance often impact a suit of other stress-related traits (Hoffmann and Parsons 1989b). Several artificial selection studies for starvation resistance have shown a correlated change in desiccation resistance, suggesting these stress traits are related (Hoffmann and Parsons 1989a; b; Chippindale *et al.* 1996; Hoffmann and Harshman 1999; Harshman *et al.* 1999; Hoffmann *et al.* 2001). For instance, a detailed study of this correlated response by Hoffmann and Parsons (1989b) demonstrated a rapid phenotypic response in both desiccation and starvation resistance following four generations of strong selection for increased desiccation resistance, and in part this was attributed to selection acting on a general stress response mechanism. Subsequent genomics studies have suggested that this rapid phenotypic response is accompanied by rapid and extensive genomic change (Kang *et al.* 2016), and that extensive pleiotropy underlies desiccation resistance (Telonis-Scott *et al.* 2012, 2016; Kang *et al.* 2016; Griffin *et al.* 2017).

We investigated the association between starvation and desiccation resistance in the DSPR by measuring female desiccation resistance in RILs chosen from the two tails of the phenotypic distribution of female starvation resistance. We found that desiccation and starvation resistance were significantly correlated (R^2^ = 43.8, F_1,31_ = 24.11, p < 0.0001; Fig 3). Since mean desiccation resistance was considerably lower than mean starvation resistance for all lines tested (compare Figs 1 and 3), flies experiencing desiccation conditions are unlikely to be dying from starvation. In addition, since DSPR lines with very low starvation resistance do not also have low larval viability (data from Marriage *et al.* 2014) or reduced adult lifespan (data from Highfill *et al.* 2016) it does not appear that DSPR lines with very low resistance to starvation and desiccation are simply “sick” (Fig S11). The relationship between starvation and desiccation resistance in the present study provides support for the genetic correlation and shared physiological mechanisms that have been proposed to exist between these traits (Hoffmann and Parsons 1989a; b, 1993; Harshman *et al.* 1999; Kennington *et al.* 2001). However, the correlation we observed is modest, and does not rule out the possibility that the covariation observed between starvation and desiccation resistance may be influenced by genetic variation in one or more other resistance-associated traits. A more intensive sampling of the DSPR would be necessary investigate the genetic correlation between starvation and desiccation resistance.

**Figure 3.**
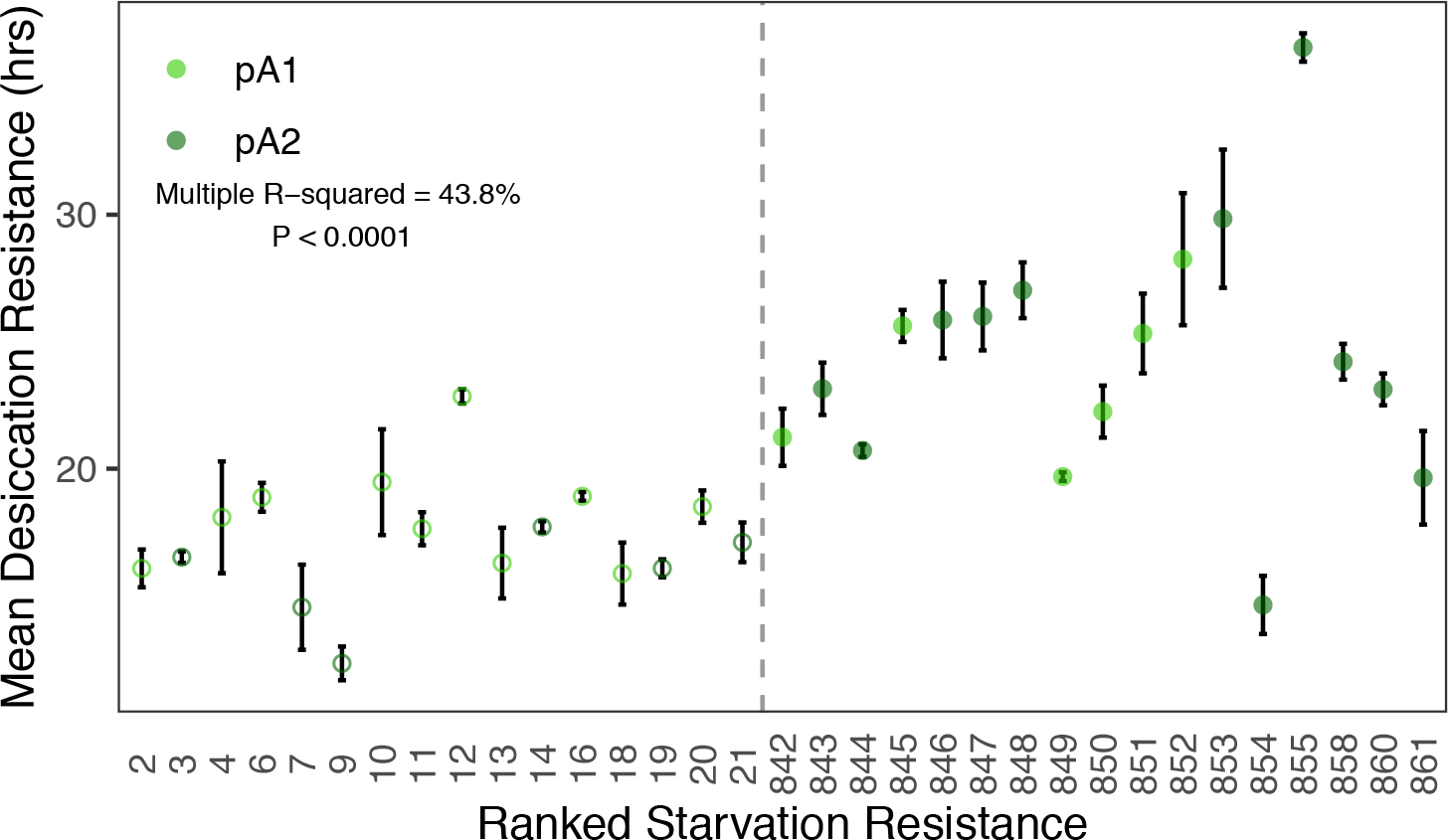
Mean starvation and desiccation resistance are correlated in the DSPR (F_1,31_ = 21.11, p < 0.0001). Desiccation resistance is presented as RIL means (±SD). Open symbols indicate “low” starvation resistance RILs; filled symbols indicate “high” starvation resistance RILs, and the dashed vertical line separates these RIL classes.

One physiological mechanism that may increase tolerance to environmental stressors is a reduction in metabolic rate (Lighton and Bartholomew 1988; Hoffmann and Parsons 1989a; b, 1991; Chippindale *et al.* 1996; Djawdan *et al.* 1997; Marron *et al.* 2003; Rion and Kawecki 2007; Schwasinger-Schmidt *et al.* 2012; Slocumb *et al.* 2015). Indeed, selection for both starvation and desiccation resistance has been shown to lead to a correlated change in activity level, an indirect proxy for metabolic rate (Hoffmann and Parsons 1989b, 1993; Schwasinger-Schmidt *et al.* 2012). Here, we assessed activity of a subset of RILs exhibiting high and low starvation resistance to understand how genetic variability in starvation resistance relates to activity levels under non-stressful conditions. In the presence of nutritive media males and females differed in activity level across the light and dark period (F_1,132_ = 16.9, p < 0.0001; Fig 4; Table S7), with high starvation resistance RILs exhibiting significantly lower activity levels than low starvation resistance RILs (F_1,132_ = 12.5, p < 0.001; Fig 4; Table S7). The effects of starvation resistance rank (high vs low), sex, and the light status (light versus dark) on activity were similar in magnitude (Cohen’s F: 0.21- 0.36; Table S7), suggesting that these factors contribute similarly to variation in waking activity levels.

**Figure 4.**
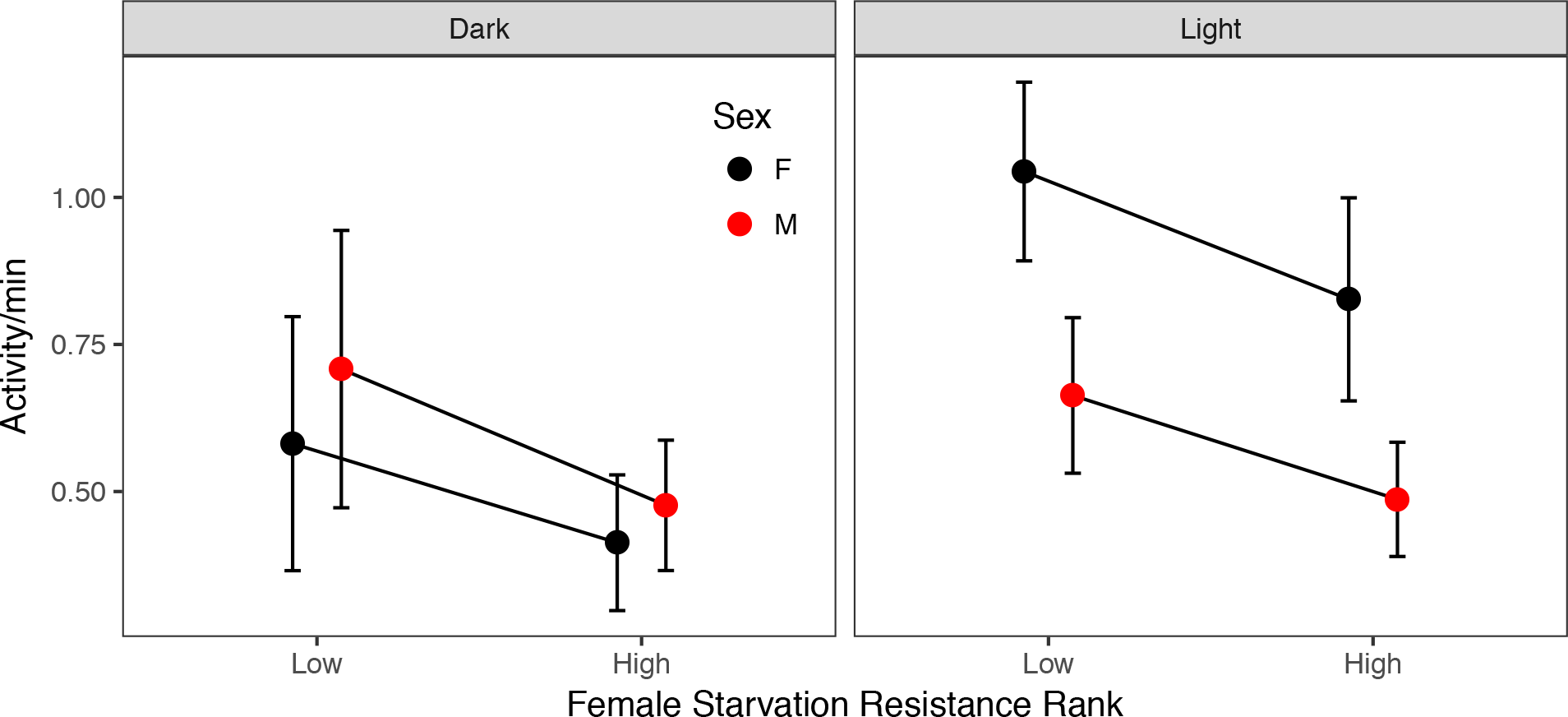
Activity level on regular media for males (red) and females (black) from a subset of high and low female starvation resistance RILs. Panels indicate the light and dark periods of a 24-hour monitoring period. Activity while awake was influenced by both a sex-by-light interaction (F_1,132_ = 16.9, P < 0.001) and by starvation resistance class (i.e. high or low; F_1,132_ = 12.5, P < 0.001).

The differences in activity between high and low starvation resistance lines on regular, nutritive media (Fig 4) were preserved under starvation stress conditions, with the high starvation resistant lines being less active than the low starvation resistant lines throughout the starvation process (Fig S13; Table S8). This pattern aligns with that from previous studies. For instance, Schwasinger-Schmidt *et al.* (2012) found that activity of flies with high starvation resistance was reduced following 15 generations of selection for starvation resistance in both males and females. Slocumb *et al.* (2015) also found that waking activity was reduced in lines selected to have high starvation resistance. Although previous associations between increased starvation tolerance and lower activity levels, metabolic rate, and changes in behavior have been observed (Murphey and Hall 1969; Hoffmann and Parsons 1989a; Blows and Hoffman 1993; Djawdan *et al.* 1997; Karan *et al.* 1998; Schwasinger-Schmidt *et al.* 2012; Masek *et al.* 2014), our findings present a novel addition to our understanding of how increased starvation resistance may occur. Behavioral components of energy conservation are likely to play a role in how individuals compensate for stressful conditions (van Dijk *et al.* 2002; McCue 2010; Masek *et al.* 2014) and represent an additional facet of the complex nature of phenotypic variability in starvation resistance.

#### Starvation resistance and triglyceride level are correlated in the DSPR

Periods of starvation have been shown to significantly reduce triglyceride levels in both males and females (Schwasinger-Schmidt *et al.* 2012), and others have suggested that fat stores and starvation resistance may be genetically correlated (Service *et al.* 1985; Rose *et al.* 1992; Chippindale *et al.* 1996; Harshman *et al.* 1999; Schwasinger-Schmidt *et al.* 2012; Slocumb *et al.* 2015). To investigate the relationship between these traits in the DSPR, we measured mean female triglyceride level in a subset of the pA and pB DSPR RILs and found substantial phenotypic and genetic variation among RILs (Table 1; Fig 5). Mean starvation resistance and triglyceride level were positively correlated in both DSPR panels, although the correlation in the pA and pB panels was significantly different (Fig 6). Overall, variation in mean starvation resistance explained 23.7% of the variation observed in mean triglyceride level among the DSPR RILs across the two mapping panels (R^2^ = 23.7%, F_3,929_ = 95.96, p < 0.0001; Fig 6), suggesting that a proportion of variation in female starvation resistance can be explained by variation in triglyceride level in the DSPR.

**Figure 5.**
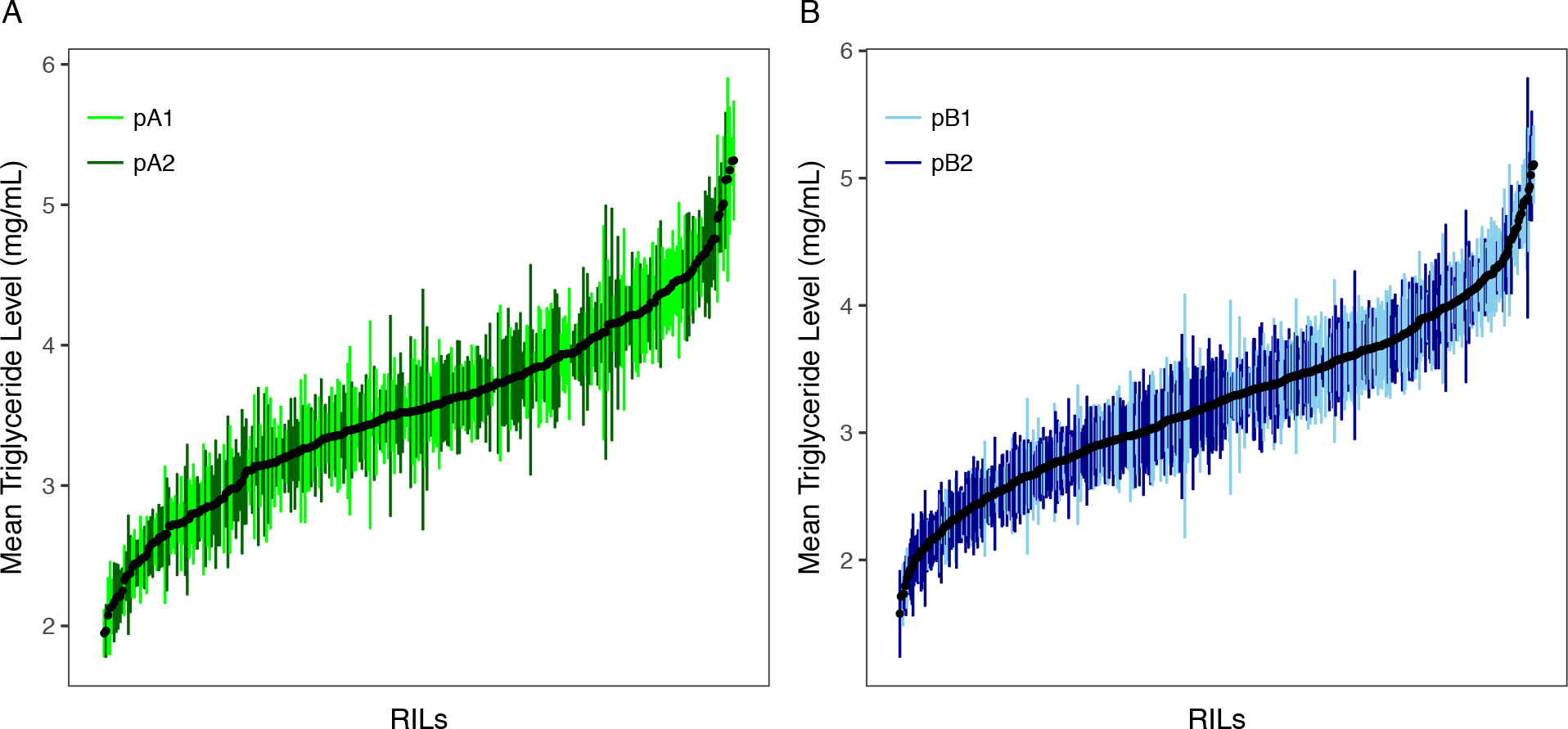
Variation in mean DSPR triglyceride level (± SD) for females in the pA panel (A) and pB panel (B).

**Figure 6.**
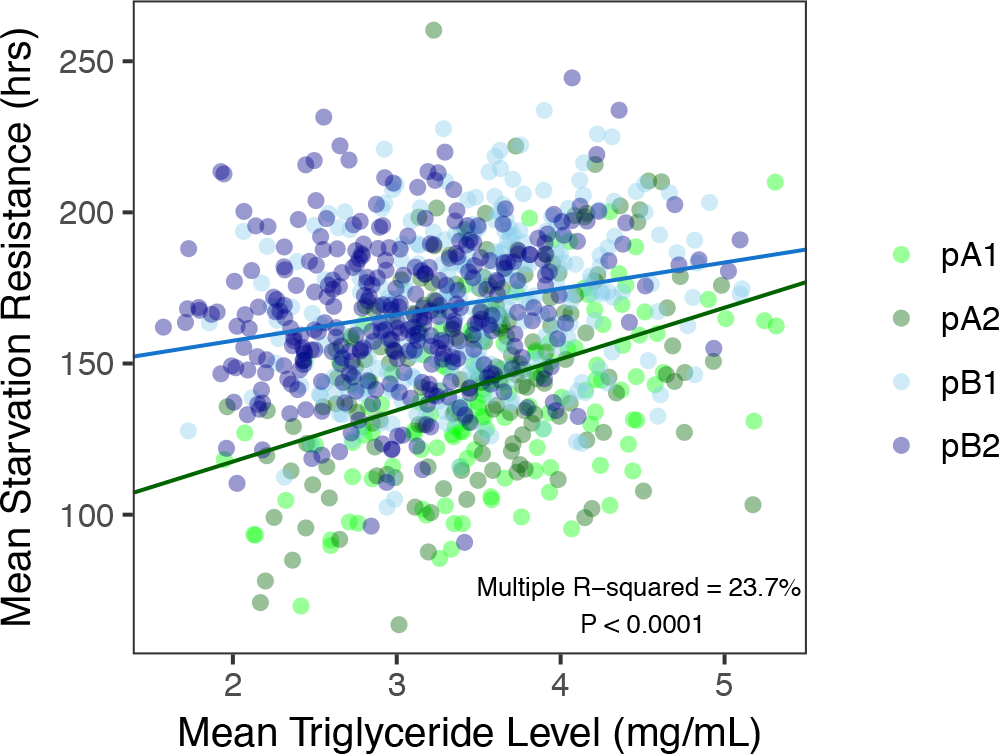
Mean starvation resistance and triglyceride level are positively correlated in females (F_3,929_ = 95.96, p < 0.0001). The strength of the correlation varied between the two mapping panels (interaction: F_1,929_ = 9.32, p < 0. 01). Points are colored to indicate subpopulation for each mapping panel, although subpopulation was not included in the regression analysis.

The correlation between triglyceride level (or total lipid level, depending on the study) and starvation resistance has been measured in numerous natural and artificially selected *Drosophila* populations, and a positive relationship is often described (Chippindale *et al.* 1996; Djawdan *et al.* 1997; van Herrewege and David 1997; Harshman *et al.* 1999; Schwasinger-Schmidt *et al.* 2012; Goenaga *et al.* 2013; Slocumb *et al.* 2015; Hardy *et al.* 2018). For example, in isofemale lines derived from populations distributed across approximately 14.4 degrees of latitude, Goenaga *et al.* (2015) found that 12% of the variation in female starvation resistance was accounted for by lipid content. Similarly, Chippindale *et al.* (1996) found a very strong positive relationship between total lipids and starvation resistance following extended selection for increased starvation resistance and suggested that lipid levels may directly determine starvation resistance. However, a strong correspondence between lipid content and starvation resistance is not always observed in strains derived from natural populations (Robinson *et al.* 2000; Hoffmann *et al.* 2001; Jumbo-Lucioni *et al.* 2010). For example, Jumbo-Lucioni *et al.* (2010) found no correlation between triacylglycerol levels and starvation resistance measured in inbred lines derived from a natural population. Hoffman *et al.* (2001) suggested that variation in the strength of the correlation between triglycerides and starvation resistance may be due to the evolutionary history of the study population. Evolutionary tradeoffs between increased lipid storage and other aspects of fitness may also influence the correlation between starvation resistance and lipid levels (Huang *et al.* 2014; Hardy *et al.* 2015, 2018). Furthermore, artificial selection may increase starvation resistance via mechanisms that preferentially modify lipid accumulation or metabolism, rather than by impacting energy level or energy-saving behavioral strategies (Hoffmann and Parsons 1989a; Blows and Hoffman 1993; Hoffmann *et al.* 2001; Marron *et al.* 2003; Rion and Kawecki 2007; Masek *et al.* 2014; Slocumb *et al.* 2015). The relationship observed between triglyceride levels and starvation resistance in our study supports the hypothesis that triglyceride levels and starvation resistance are likely physiologically related. Equally, it is evident from our data that triglyceride level likely influences starvation resistance to a lesser degree than proposed by Chippindale *et al.* (1996) and Hoffmann and Harshman (1999), and that starvation resistance and triglyceride level have the potential to evolve independently under natural selection.

#### Starvation resistance QTL allow prediction of DSPR founder phenotypes

We identified 8 QTL for starvation resistance in the pA panel and 7 QTL in the pB panel, several of which overlapped between sexes (Fig S14). Both sets of QTL explained a substantial amount of variation in starvation resistance, with individual peaks accounting for 3.7-13.2% of the variation (Table 2). The total variance explained by QTL in the pA (pB) population was 26.1% (32.8%) in females and 17.5% (37.9%) in males, assuming QTL are independent and additive (Table 2). None of the QTL identified in the pA and pB mapping panels overlapped, and since power to detect 5% QTL is expected to be high in our study (King *et al.* 2012a) and all DSPR phenotyping was completed within seven months using the same design and environmental conditions, this likely reflects genetic variation among the different sets of founders used to establish the two sets of lines.

**Table 2.** Summary of QTL identified for starvation resistance and triglyceride level in the DSPR.

| Starvation Resistance QTL: pA DSPR panel |  |  |  |  |  |  |  |  |
| --- | --- | --- | --- | --- | --- | --- | --- | --- |
| QTL | Sex | Peak LOD | Chr | Physical Interval (Mb) | Genetic Interval (cM) | Variance Explained | No. Genes | No. Potential Starvation Candidate Genes<sup>a</sup> |
| SA1 | M | 7.41 | 2L | 0.34-0.64 | 0.36-1.02 | 3.90 | 52 | 3 |
| SA2 | F | 10.85 | 2R | 4.90-5.53 | 59.39-60.64 | 6.01 | 122 | 10 |
| SA2 | M | 10.59 | 2R | 4.87-5.58 | 59.32-60.74 | 5.52 | 126 | 10 |
| SA3 | F | 8.05 | 2R | 12.85-13.43 | 80.75-82.81 | 4.23 | 110 | 9 |
| SA4 | F | 7.03 | 2R | 14.18-14.72 | 84.14-86.68 | 3.70 | 105 | 7 |
| SA5 | F | 10.13 | 3L | 3.40-3.92 | 6.89-8.88 | 5.29 | 52 | 4 |
| SA6 | F | 13.21 | 3R | 0.79-2.37 | 47.10-47.52 | 6.85 | 244 | 11 |
| SA7 | M | 7.66 | 3R | 15.55-15.86 | 66.52-67.36 | 4.03 | 52 | 4 |
| SA8 | M | 7.73 | 3R | 21.07-21.48 | 86.77-88.20 | 4.06 | 80 | 3 |
| Starvation Resistance QTL: pB DSPR panel |  |  |  |  |  |  |  |  |
| QTL | Sex | Peak LOD | Chr | Physical Interval (Mb) | Genetic Interval (cM) | Variance Explained | No. Genes | No. Potential Starvation Candidate Genes<sup>a</sup> |
| SB1 | F | 7.68 | 2L | 7.02-7.30 | 23.69-25.20 | 4.05 | 33 | 1 |
| SB2 | M | 8.35 | 2L | 10.10-10.50 | 39.24-40.93 | 4.39 | 110 | 13 |
| SB3 | F | 7.13 | 2L | 12.14-12.65 | 45.94-46.96 | 3.77 | 44 | 4 |
| SB3 | M | 7.05 | 2L | 12.13-12.63 | 45.92-46.93 | 3.72 | 45 | 4 |
| SB4 | F | 12.28 | 2R | 1.10-2.70 | 54.93-55.73 | 6.41 | 148 | 10 |
| SB4 | M | 19.40 | 2R | 2.01-2.69 | 55.21-55.72 | 9.91 | 81 | 8 |
| SB5 | F | 18.98 | 3L | 18.94-19.42 | 45.48-45.75 | 9.73 | 54 | 5 |
| SB5 | M | 26.40 | 3L | 19.00-19.43 | 45.51-45.76 | 13.24 | 46 | 3 |
| SB6 | F | 9.07 | 3R | 8.86-9.20 | 52.50-53.29 | 4.77 | 48 | 7 |
| SB6 | M | 12.77 | 3R | 8.57-9.21 | 52.33-53.30 | 6.64 | 90 | 9 |
| SB7 | F | 7.72 | 3R | 25.65-26.15 | 99.12-100.11 | 4.08 | 111 | 8 |

**Triglyceride Level QTL: pB DSPR panel**
| QTL | Sex | Peak LOD | Chr | Physical Interval (Mb) | Genetic Interval (cM) | Variance Explained | No. Genes | No. Potential Starvation Candidate Genes<sup>a</sup> |
| --- | --- | --- | --- | --- | --- | --- | --- | --- |
| TB1 | F | 8.73 | 3L | 3.45-4.09 | 7.08-9.52 | 6.22 | 84 | 7 |
| TB2 | F | 7.90 | 3R | 7.18-8.01 | 50.50-51.54 | 5.64 | 118 | 5 |
| TB3 | F | 8.45 | 3R | 8.31-8.71 | 51.96-52.54 | 6.03 | 54 | 8 |
| TB4 | F | 7.71 | X | 17.34-17.43 | 57.17-57.44 | 5.51 | 4 | 0 |

<sup>a</sup>Genes and functions associated with starvation resistance are listed in Table S9. The number of genes associated with starvation resistance (SR) was determined by cross-referencing genes within each QTL interval with previously reported candidates and biological functions reported by FlyBase (Lin et al. 1998; Clancy et al. 2001; FlyBase Curators et al. 2004; Harbison et al. 2004; Sørensen et al. 2007; Nuzhdin et al. 2007).

Because we can estimate the effect associated with each founder haplotype at each mapped QTL in the DSPR, it follows that a combination of the estimates across all QTL can be used to predict the actual phenotypes of the original founder strains. We measured starvation resistance in the 15 DSPR founder lines (Fig 1E, F) to test this prediction. It is likely that the strength of the correlation between the estimated and actual trait response is influenced by the number and effect size of each QTL mapped for the trait. To account for differences in the degree to which starvation resistance is influenced by QTL of varying effect sizes, we calculated the predicted mean trait for each founder line weighted by the variance explained by each QTL.

As anticipated, using a general linear model the weighted mean predicted starvation resistance of the founders based on QTL effects was significantly correlated with the sex-averaged mean starvation resistance measured for the founder lines (R^2^ = 60.8%, F_3,13_ = 6.12, p < 0.01; Fig 7). The slope of this relationship is relatively small (β = 0.13 ± 0.07), suggesting that while a large component of variation in starvation resistance is clearly genetic (supported by heritability estimates for each panel, Table 1), substantial variation in the phenotype is unaccounted for by additive genetic effects at mapped QTL. This unaccounted-for genetic variation in starvation resistance is likely due to many QTL with very small effects beyond our power to detect them (King *et al.* 2012a) and/or epistatic interactions among QTL (Evans *et al.*; Mackay 2014). Epistasis may be especially important when comparing actual founder strain phenotypes with those inferred via QTL effects due to the many generations of recombination employed while establishing the DSPR from the inbred founders. However, the strength of the correlation between predicted and actual responses does suggest that QTL identified from the DSPR mapping panels identify causative loci that influence the level of starvation resistance among the progenitors of the RILs.

**Figure 7.**
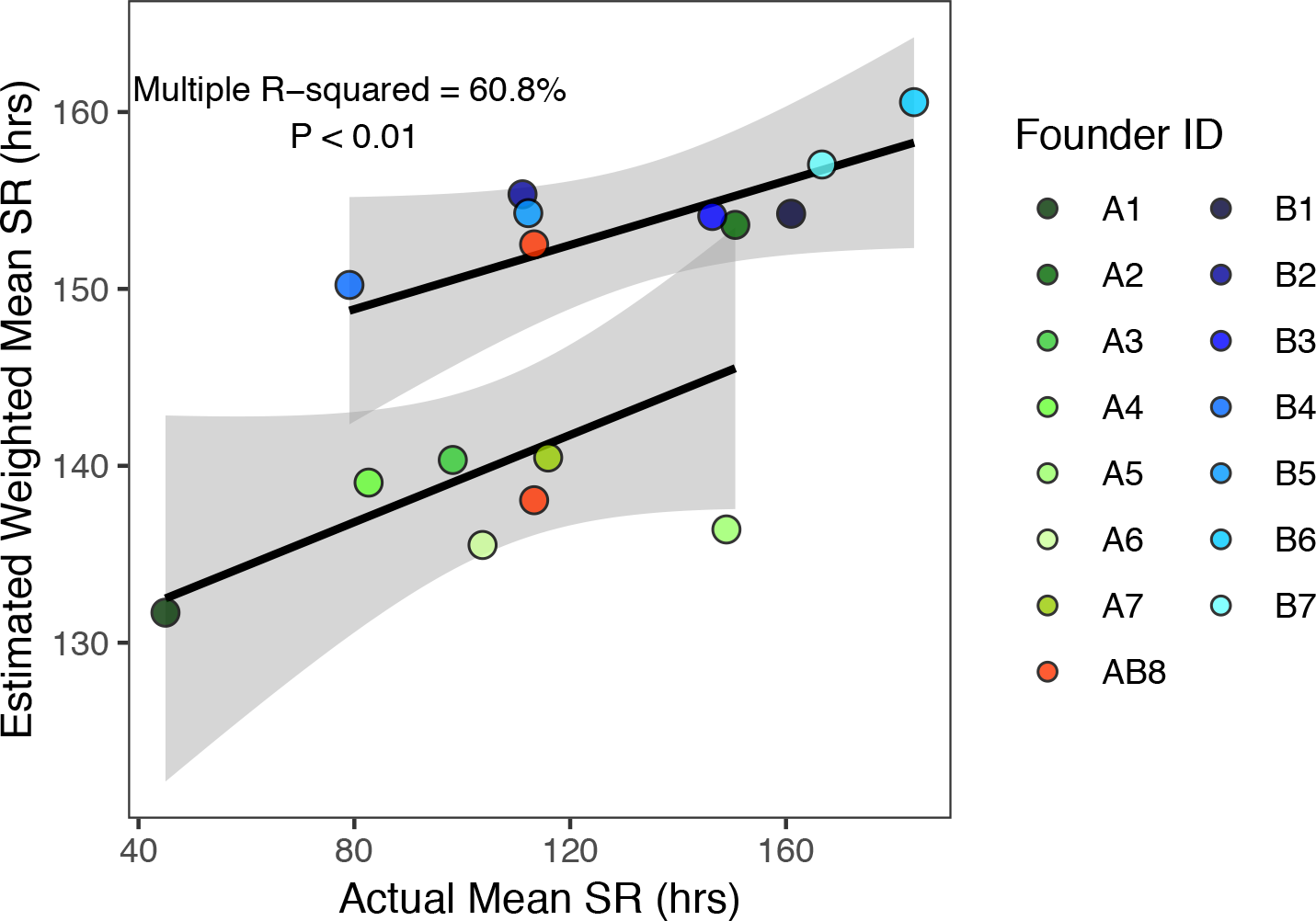
Estimated starvation resistance weighted by the variance explained by each QTL and actual starvation resistance measured for the 15 founder lines of the DSPR were significantly correlated (β = 0.13±0 0.07; F_3,13_ = 6.21, p < 0.01). AB8 identifies the founder line shared between the pA and pB mapping panels estimated independently in each QTL analysis. Grey shading indicates the 95% CI of the regression.

#### Limited overlap between the genetic architecture of starvation resistance and triglyceride level in the DSPR

To further understand the relationship between starvation resistance and triglyceride level in the DSPR we compared the genetic architectures of these two traits. We mapped four distinct QTL for triglyceride level in the pB population, each of which accounted for 5.5-6.2% of the variation in this trait (Table 2; Fig S15), in total explaining 23.4% of the variation in pB. No QTL for triglyceride level were detected in the pA panel, likely due to the reduced number of pA RILs assessed (pA N = 311; pB N = 628). However, even with this reduced power, the QTL map for pA suggests that the genetic architecture for triglyceride level is different between the two mapping panels, as there is no evidence of near-significant peaks in pA within QTL intervals statistically identified in the pB panel (Fig S15).

Given the phenotypic correlation between triglyceride level and starvation resistance in the DSPR (Fig 6) and similar correlations previously reported in other studies (Chippindale *et al.* 1998; Hoffmann and Harshman 1999), one might predict overlap of QTL associated with these traits. However, we see only limited evidence for this. Triglyceride QTL TB1 (mapped in the pB panel) and starvation QTL SA5 (mapped in the pA panel) do physically overlap, but given the complete lack of evidence for QTL for the same trait co-localizing in both the pA and pB DSPR mapping panels, it is unlikely the variant(s) underlying these QTL are the same. To investigate the relationship between the two QTL that do overlap within the same panel (SB6 and TB3), we assessed the influence of haplotype structure at the overlapping QTL on the positive phenotypic correlation between triglyceride level and starvation resistance (Fig 6). In this analysis, we first identified the founder haplotype for each RIL at the positions of the overlapping QTL peaks, and calculated the average phenotype of each of the founder haplotypes. We then assessed the correlation between haplotype-specific mean triglyceride level and starvation resistance with a general linear model. After accounting for the haplotype structure at the overlapping peaks, we found that mean starvation resistance and triglyceride level were significantly correlated (F_1,7_ = 7.72, p < 0.05, R^2 =^ 52.4%; Fig S16), suggesting some pleiotropic variants may be responsible for this pair of overlapping starvation resistance and triglyceride level QTL.

The limited overlap in the QTL intervals associated with starvation and triglyceride level suggests that the genetic bases of this pair of traits are largely independent, or at least not tightly linked at QTL with moderate to large effects. In natural populations, increased starvation resistance may evolve as a result of selection on diverse traits including metabolic rate, activity level, lifespan, development rate, thermal tolerance, and fecundity (Service *et al.* 1985; Hoffmann and Parsons 1989b, 1993; Rose *et al.* 1992; Chippindale *et al.* 1993; Djawdan *et al.* 1997; Harshman *et al.* 1999; Bochdanovits and de Jong 2003; Marron *et al.* 2003; Bubliy and Loeschcke 2005; Rion and Kawecki 2007; Schwasinger-Schmidt *et al.* 2012), and triglyceride levels may be influenced by genetic variation in each of these traits. Our evidence of minimal overlap between the genetic architectures of starvation resistance and triglyceride levels, coupled with a phenotypic correlation between these traits, may be indicative of a series of complex correlations between traits that influence stress tolerance, energy metabolism, and life history in the DSPR.

#### Candidate genes underlying fitness trait variation

Across all QTL identified for starvation resistance and triglyceride level in this study, several genes within mapped QTL intervals have functions related to these and other correlated traits (Table 2, Table S9). Of particular interest are the 30 genes that fall within our QTL intervals that were identified in previous starvation resistance studies (Clancy *et al.* 2001; Harbison *et al.* 2005; Sørensen *et al.* 2007) (Table S9). Gene ontology analyses performed for each trait and panel revealed enrichment of genes within pA starvation QTL related to glutathione metabolic process (6.91-fold enrichment, FDR corrected P = 0.000146; Table S10), as well as several categories that were enriched for genes implicated by mapped triglyceride QTL (Table S10). This enrichment could assist with the resolution of the functional genes within QTL regions. However, it should be noted that the sets of genes implicated by QTL mapping in the DSPR (3-244 genes per interval in this study) are extremely unlikely to all contribute to trait variation, and their presence within QTL intervals cannot alone be taken as evidence for causality.

Upon examination of the genes within the overlapping starvation resistance and triglyceride interval (TB3 and SB6), we found several genes that have either been predicted or experimentally demonstrated to be associated with traits related to starvation resistance and triglycerides or metabolism (Table S9 and references therein). Genes that fall within the intervals of the overlapping peaks include those that influence adult lifespan (e.g. *ry*, *Men*, *Gnmt* (Simonsen *et al.* 2006; Paik *et al.* 2012; Obata and Miura 2015)), lipid metabolic processes (including *Lip3*, *CG11598*, *CG11608*, *CG18530* (FlyBase Curators *et al.* 2004)), insulin signaling (e.g. *poly* (Bolukbasi *et al.* 2012)), response to starvation (e.g. *mthl12*, *Gnmt* (UniProt Curators 2002; Obata *et al.* 2014)), larval feeding behavior (e.g. *Hug* (Melcher and Pankratz 2005)), circadian rhythm and sleep (e.g. *timeout*, *Men* (Harbison *et al.* 2004; Benna *et al.* 2010)), and triglyceride homeostasis (*Gnmt* (Obata *et al.* 2014)) (Table S9). These genes are promising candidates for future studies seeking to examine the functional genetic relationship between these two traits.

#### Different mapping approaches reveal unique genetic architectures for starvation resistance

Dissection of a quantitative genetic trait using different approaches can allow greater resolution of the genetic architecture, and provide insight into how alleles unique to different mapping panels contribute to phenotypic variation. To gain this additional understanding we assessed the genetic architecture of starvation resistance in the DGRP using GWA mapping of four starvation resistance datasets: new data collected in this study, data from Mackay *et al.* (2012), data from Everman and Morgan (2018), and a consensus, across-study starvation resistance measure calculated as the mean response across the three starvation datasets (150 lines were measured across the three studies). Using an FDR threshold of 20%, between 0 and 12 SNPs were associated with starvation resistance in each dataset and sex (Table 3; Table S11). Aside from 3 SNPs that overlap between the across-study mean dataset and the Mackay *et al.* (2012) dataset, none of these above-threshold SNPs were the same (Table 3; Table S11). Using the more lenient significance threshold of P < 10^−5^ (see Table 3 for the equivalent FDR values), between 17 and 48 SNPs were associated with starvation resistance for each dataset and sex (Table 3). However, overlap in associated SNPs among datasets was still minimal (Fig S17). The SNPs identified using the more lenient significance threshold include all those identified at the FDR threshold of 20%, so all subsequent analyses are performed on the larger set of associated SNPs, and we acknowledge that these sets may include larger fractions of false-positive associations.

**Table 3.** Summary of DGRP GWA results and lines measured in this study, Mackay et al. (2012) and Everman and Morgan (2018).

| Study<sup>a</sup> | Significance Threshold | Equivalent FDR |  | No. Strains | No. SNPs<sup>b</sup>: |  |
| --- | --- | --- | --- | --- | --- | --- |
|  |  | Female | Male |  | Female | Male |
| This Study | FDR 0.2 | - | - | 168 | 2 | 3 |
| | $P < 10^{-5}$ | 0.78 | 0.79 | 168 | 23 | 19 |
| *Mackay et al. (2012)* | *FDR 0.2* | - | - | 203 | 11 | 0 |
|  | *$P < 10^{-5}$* | 0.49 | 0.82 | 203 | 39* | 17* |
| *Everman and Morgan (2018)* | *FDR 0.2* | - | - | 164 | 0 | 0 |
|  | *$P < 10^{-5}$* | 0.67 | 0.65 | 164 | 25* | 22* |
| Across-Study Mean Response | FDR 0.2 | - | - | 150 | 12 | 0 |
| | $P < 10^{-5}$ | 0.43 | 0.80 | 150 | 48 | 22 |

<sup>a</sup>Text in italics indicates results from the reanalysis of starvation data previously presented by Mackay et al. (2012) and Huang et al. (2014) and Everman and Morgan (2018).
<sup>b</sup>Asterisks indicate results that match those reported in the original studies.

Across all four datasets reporting starvation resistance in the DGRP, 12 SNPs (associated with seven genes) identified using the P < 10^−5^ significance threshold fell within QTL intervals identified for starvation resistance in the DSPR (Fig S18; Table S11). In females, one SNP associated with starvation resistance from Mackay *et al.* (2012) is within QTL SA4 (gene *CG30118*), and one SNP associated with starvation resistance in this study is present in QTL SA3 (gene *mbl*). One SNP associated with starvation resistance from Everman and Morgan (2018) is within QTL SB7 (gene *hdc*), and one SNP from the average of starvation resistance across the DGRP datasets is within QTL SB5 (gene *Gbs-76A*). In males, one SNP associated with starvation resistance from Mackay *et al.* (2012) is within QTL SB3 and was not associated with a gene. Four SNPs (one from this study and three from the average of starvation resistance) fell within QTL SB5 and were all near the gene *fz2*; one additional SNP associated with the average of starvation resistance was also within SB5 and was associated with the gene *pip*. One SNP associated with starvation resistance from Everman and Morgan (2018) was within QTL SB6 (gene *beat-Vc*). Only one of these seven genes has been previously associated with starvation resistance (*CG30118*; (Sørensen *et al.* 2007)), and none have reported functions specifically related to starvation resistance or general stress response (FlyBase Curators *et al.* 2004). Furthermore, none of the overlapping genes survived an FDR threshold of 0.2, increasing the possibility that these genes may be false positives. Therefore, with the possible exception of *CG30118*, these genes may not be promising candidates, despite their overlap among studies.

Compared to genes implicated by QTL identified in the DSPR, which include several that have been previously associated with starvation or related phenotypes (e.g., lifespan or lipid content), DGRP GWAS hits implicate fewer *a priori* strong candidate genes. Additionally, we did not observe any GO enrichment following analyses of SNPs associated with the four starvation resistance datasets, although we acknowledge that the limited number of implicated genes likely compromised the power of these analyses. Of the total 127 unique genes associated with starvation resistance in the DGRP across studies and sexes, only two have been previously identified as associated with starvation resistance in other mapping populations (*CG30118*, *scaf6*; Table S11; (see Table S2 in Sørensen *et al.* 2007)). More generally, five had previously been associated with the determination of adult lifespan (e.g. *cnc* and *Egfr*; Table S11; (Sykiotis and Bohmann 2008; Kamakura 2011)), and 5 have been previously associated with lipid metabolism or metabolic processes (e.g. *GlcAT-P* and *Ugt86Dj*; Table S11; (FlyBase Curators *et al.* 2004; Gaudet *et al.* 2010)). Given the relative lack of power of a GWA study using less than 200 genotypes (Long *et al.* 2014), and our use of a permissive genomewide threshold, it could be that many of the GWAS associations are incorrect, explaining why associations do not typically tag known candidates. Equally, it could be the case that a series of novel pathways are involved in natural variation for starvation resistance, and that traditional candidates - often identified via mutagenesis screens rather than through examination of segregating allelic variation - typically do not harbor the functional natural variants detectable in a GWAS (Mackay *et al.* 2009).

The general lack of correspondence among the loci associated with starvation resistance in each mapping panel does not invalidate either approach as a strategy to uncover functional variation. It is likely that many genes contribute to variation in this trait with effects that are either fairly small, or that have effects only in a specific genetic background (i.e., exhibit genetic epistasis), and we would not expect to routinely identify such loci. In addition, comparison of the genetic architecture of quantitative genetic traits across multiple panels is complicated by a number of additional factors. The genetic structure of the mapping panel (e.g., whether it is a multiparental panel like the DSPR or a population-based association study panel like the DGRP) influences the analytical strategy, and the power and resolution of mapping (Long et al. 2014). The complement of alleles present in the panel, and the frequency with which they segregate, will also affect the ability to identify the same locus across mapping panels (e.g., King and Long 2017). This point is especially true for the comparisons made here, since the DSPR represents a global sampling of genetic variation represented by the 15 founder strains, whereas the variation present in the DGRP is a direct reflection of the genetic variability in a single population at a single point in time. Therefore, a lack of overlap in the identified QTL for a complex, highly-polygenic trait between the DSPR and the DGRP is perhaps not unexpected.

#### Repeatability in the SNPs associated with starvation resistance across DGRP studies

The public availability of starvation resistance data for the DGRP from multiple studies provides a novel opportunity to investigate the reliability and repeatability of associations identified for a classic quantitative trait in the same mapping panel across independent phenotypic screens (Lithgow *et al.* 2017). Despite the moderately high phenotypic correlation between starvation resistance measured in the three studies (Fig S8), only a single variant was implicated in more than one of the three studies (Fig S17).

The lack of overlap in SNPs associated with starvation resistance could be due to differences in the rearing/testing environments of the three studies (discussed above), where genotype-by-environment effects - often pervasive for complex traits (Gurganus *et al.* 1998) - could lead to different sites impacting variation in different studies. However, it is potentially more likely that the sets of associated SNPs have real, but extremely small effects on starvation resistance variation, and power in a GWA panel of modest size is too low to consistently detect them (Boyle *et al.* 2017). If true, one would predict that - in contrast to random SNPs - the “top SNPs” identified within each starvation resistance dataset would have additive effects of the same sign across all studies. In essence, significantly associated SNPs with positive effects on starvation resistance from data collected in this study would be expected to have positive additive effects on starvation resistance measured in Mackay *et al.* (2012) and Everman and Morgan (2018) more often than expected by random chance.

To test this prediction, we first collected the sign of the additive effects of SNPs that survived the significance threshold (P < 10^−5^) in each dataset for both sexes (Table S11), and determined whether these top SNPs had additive effects of the same sign in every other dataset. We then established a null distribution of SNP additive effect signs across pairs of datasets. This was accomplished by taking samples of 50 SNPs segregating in the DGRP and extracting the sign of the additive effect of each SNP in the pair of datasets, regardless of the association statistic for that SNP. The proportion of the 50 SNPs that had a positive additive effect on the trait was recorded for each of 1000 iterations and used to build an expected distribution of SNP effect sign sharing for each pair of datasets. We note that to compensate for any allele frequency bias in the variants that are actually most associated with phenotype, we ensured that the frequencies of the randomly-sampled SNPs were stratified to match the top 50 SNPs associated with each sex and dataset (see Materials and Methods; Table S3).

Finally, we compared the proportion of top SNPs for which the sign stayed the same across studies to our expected distribution for each pair of datasets (Fig 8). We found that for the random samples of SNPs, the probability that the additive effects were positive in both datasets compared was greater than 50% for most comparisons (distribution means ranging from 45.9- 72.1%; Fig 8). This implies that a random SNP is slightly more likely to have the same sign effect across data sets, which may be explained in part by the phenotypic correlation between the datasets (Fig S10). Even given this skew, far more top SNPs than expected by chance had additive effects of the same sign in each of the other starvation resistance studies (Fig 8; Table S3; Fig S19). This may suggest that there is phenotypic signal even in SNPs that have very small effects, and that are not clearly associated with starvation resistance in the GWAS (see Yang *et al.* 2010). Generally, the consistency of the additive effects of SNPs associated with starvation resistance in the DGRP calculated across datasets implies that starvation resistance is a highly polygenic trait, with a large number of QTL with very small effects that influence variation in this trait (Mackay 2004; Boyle *et al.* 2017).

**Figure 8.**
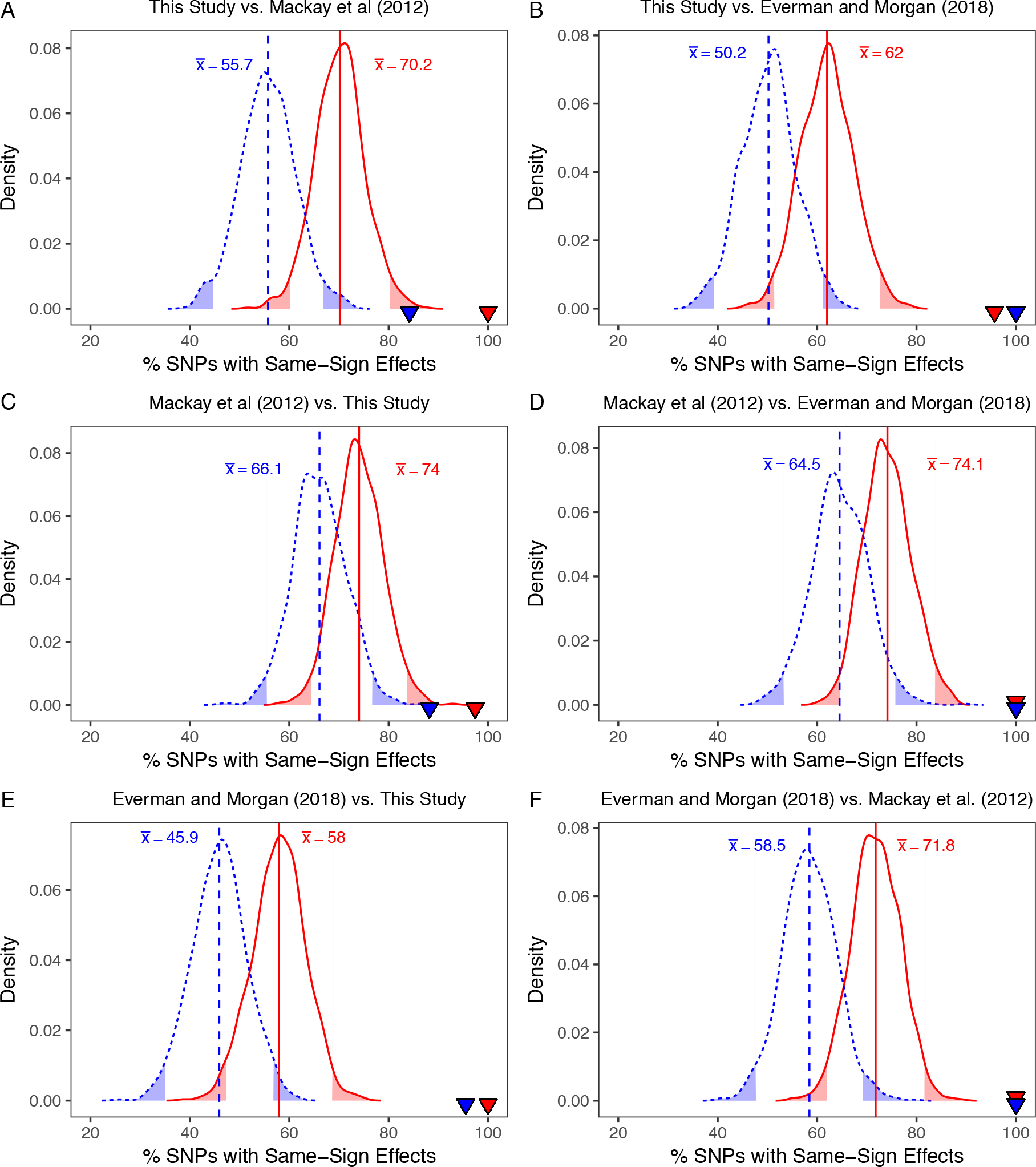
Distribution of the proportion of SNPs that are expected to have additive effects of the same sign in pairwise comparisons of data from this study, Mackay et al. (2012) and Everman and Morgan (2018). Data shown were obtained for random samples of 50 SNPs with 1000 iterations. In each plot, red indicates female data, blue indicates male data, and corresponding vertical lines and text annotation indicate the mean of the sex-specific samples. The shaded tails represent the upper and lower 95% confidence intervals of each distribution. The triangles in each plot represent the sex-specific observed proportion of top SNPs from each GWA analysis (P < 10^−5^) that had additive effects that were the same sign across studies (Fig S19). For example, in A, 100% of the SNPs associated with female starvation resistance in this study had additive effects of the same sign when calculated for starvation resistance reported in Mackay et al. (2012). The dataset comparison is indicated above each plot. In every case, top SNPs from one study were more likely to have the same additive effect sign in a second study than a random set of frequency-matched SNPs.

## Conclusions

In this study, we have described the complex quantitative genetics of starvation resistance in two large *D. melanogaster* mapping panels that have been thoroughly genomically characterized. The DSPR and DGRP panels have the advantage of increased, stable genetic diversity and provide a unique comparison to many previous quantitative genetic studies of starvation resistance that may be limited by genetic diversity or mapping power. Correlations between starvation resistance and the additional traits described in this study offer insight into the genetic control of related stress response traits and provide support for the hypothesis that the genetic architecture of stress traits varies by population and is dependent upon sex, environment, and the evolutionary history of the populations studied. The relationships between the traits analyzed in this study also offer insight into the broader responses of organisms to starvation stress, given the high conservation of mechanisms related to starvation resistance in diverse species (Partridge *et al.* 2005; Rion and Kawecki 2007). Here, we have demonstrated that traits related to survival under starvation conditions, energy storage, activity levels, and survival under desiccation stress are phenotypically correlated in the DSPR, consistent with previous artificial selection studies as well as some natural populations. However, we also clearly demonstrate that starvation resistance and triglyceride level are largely genetically independent traits, indicating that evolutionary constraint between these two traits is unlikely. We additionally describe the highly polygenic nature of starvation resistance using the DGRP, leveraging previously published phenotypes on the same lines to compare the genetic architecture of the trait across three studies. Our work shows that despite a lack of overlap across studies in the identity of the variants associated with phenotype at a nominal genomewide threshold, the sign of the additive effects of such top SNPs are conserved across studies conducted by different labs. In turn this suggests that these variants do contribute to the phenotype but have sufficiently small effects that they are not routinely captured following a severe, genomewide correction for multiple tests. From this, we gain a much more detailed understanding of the genetic control of trait variation in a genetically diverse panel and provide insight into the utility of across-study and across-panel comparisons.

## Acknowledgments

We thank Sophia Loschky and Brittny Smith for assistance with data collection, Tony Long and Libby King for assistance with analysis of early versions of the datasets reported here, and both John Kelly and Libby King for comments on a previous version of this manuscript. We also thank two anonymous reviewers and Corbin Jones for their constructive input on an earlier version of this manuscript. The DGRP collection was purchased from the Bloomington Drosophila Stock Center (NIH P40 OD018537). The project was supported by NIH grant R01 OD010974 to S.J.M. and Anthony D. Long. E.R.E. was partially supported by a Postdoctoral Fellowship from the Kansas INBRE (P20 GM103418).

## Supplemental Figures

**Figure S1.**
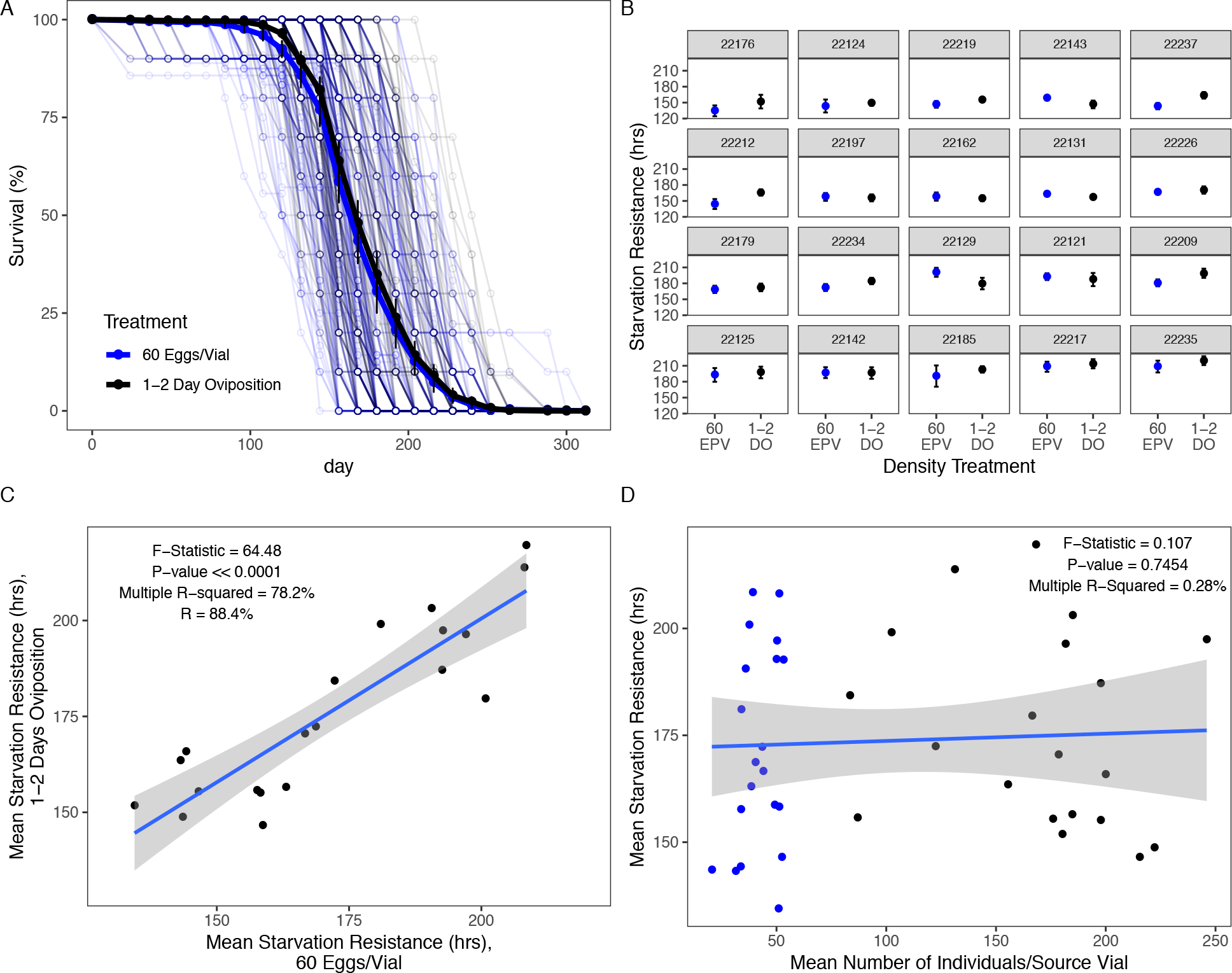
Egg density in vials used to generate experimental individuals has a limited influence on starvation resistance. This was tested in 20 randomly-selected DSPR RILs by rearing experimental individuals according to two treatments: with 60 eggs per vial (60 EPV) or with 1-2 days of oviposition (1- 2 DO). The total number of adults emerging from each source vial was also counted. Starvation resistance of the experimental individuals from each density treatment was measured as described for the large-scale starvation screen. A. Percent survival per vial at each 12-hr assessment point was very similar throughout the course of this experiment regardless of rearing density. Bold lines and points indicate the overall mean (±95% CI) survival for each treatment group at each 12-hr assessment point. B. Mean (±95% CI) starvation resistance for each of the 20 randomly-selected DSPR RILs was rarely influenced by the density treatment. Overall, density treatment had a minor effect on the average lifespan of each DSPR RIL (F_1,19_ = 18.15, P < 0.0001, % Variance Explained = 0.90%; see Table S1 for full breakdown of variance components). C. Mean starvation resistance by DSPR RIL was strongly correlated between the two density treatments. D. The mean number of individuals per source vial of experimental flies did not explain a significant amount of variation in starvation resistance. In A, B, and D, black corresponds to the 1-2 day oviposition treatment; blue corresponds to the 60 eggs per vial treatment. In C and D, grey shading represents the 95% CI of the regression.

**Figure S2.**
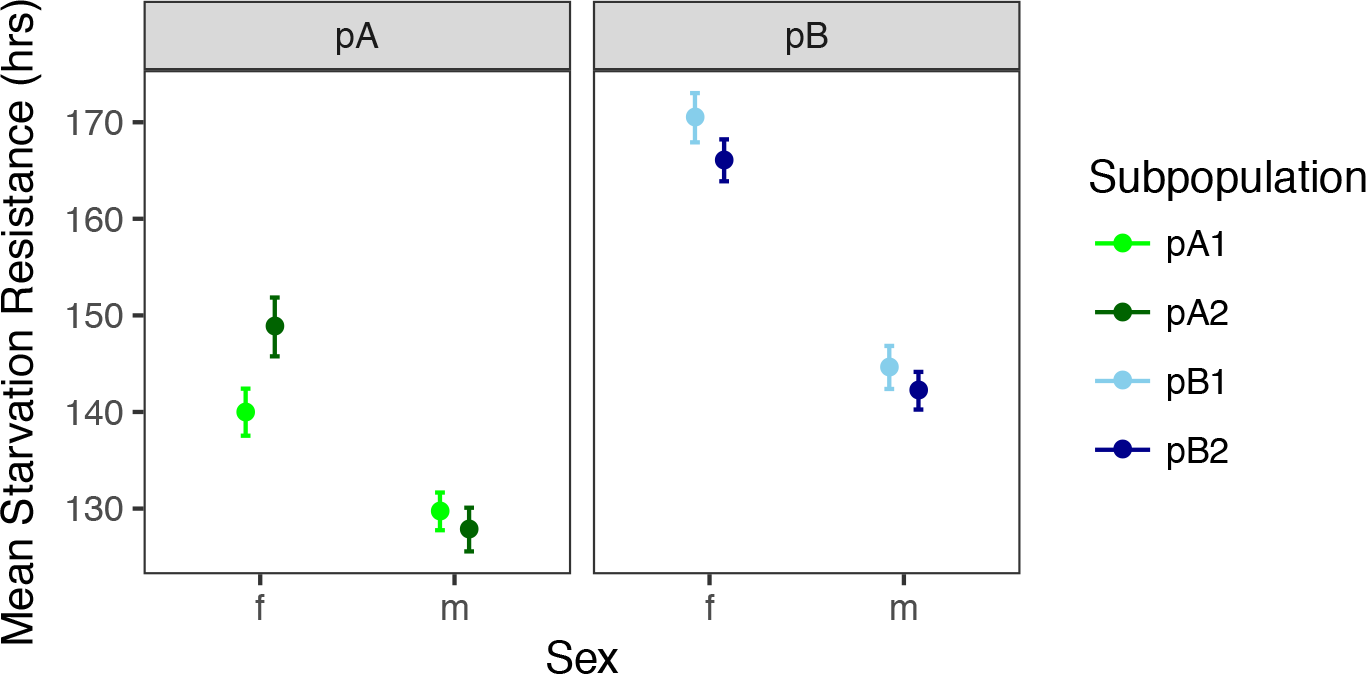
Mean (M 95% CI) starvation resistance per subpopulation and sex. Sex and subpopulation interact to influence starvation resistance (F_3,3440_ = 18.317; p < 0.0001), though only females from the pA1 and pA2 subpopulations were significantly different within a panel (Tukey’s HSD adj. p < 0.0001).

**Figure S3.**
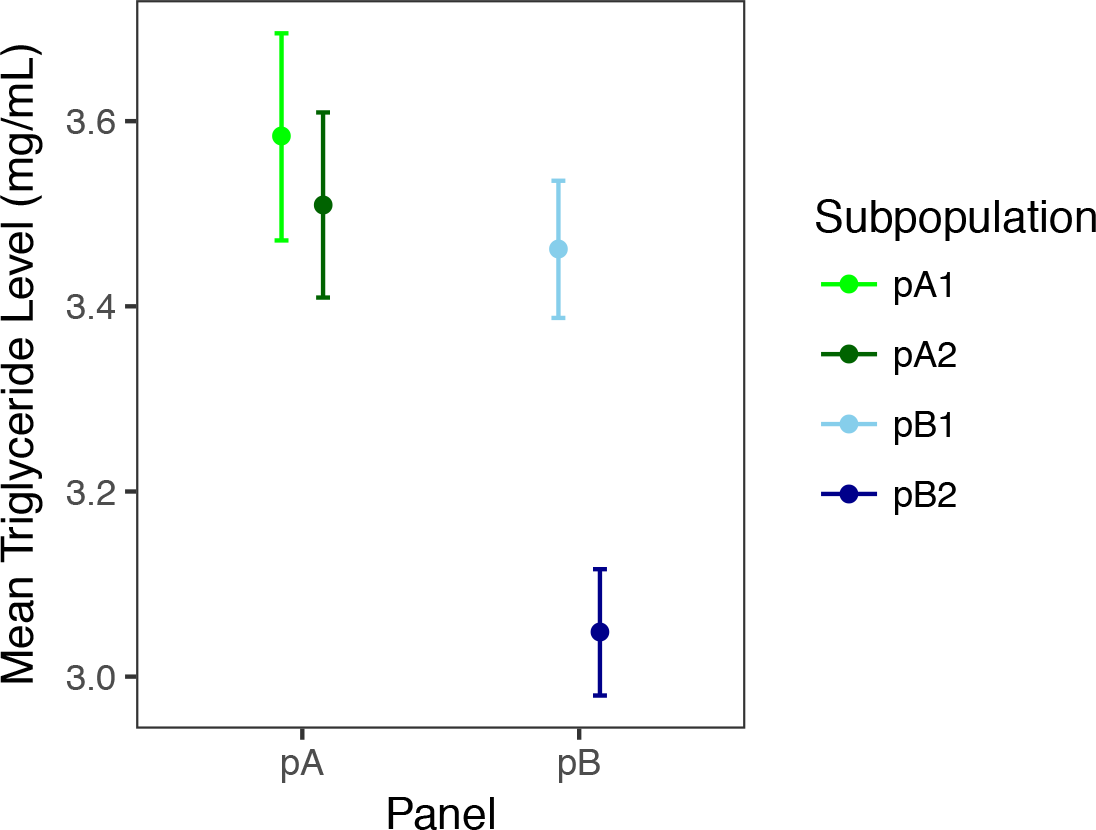
Mean female triglyceride level per subpopulation (±95% CI). Subpopulation influenced triglyceride level (F_3,935_ = 37.099; p < 0.0001), though this was driven by differences between the pB subpopulations (Tukey’s HSD adj. p < 0.0001)

**Figure S4.**
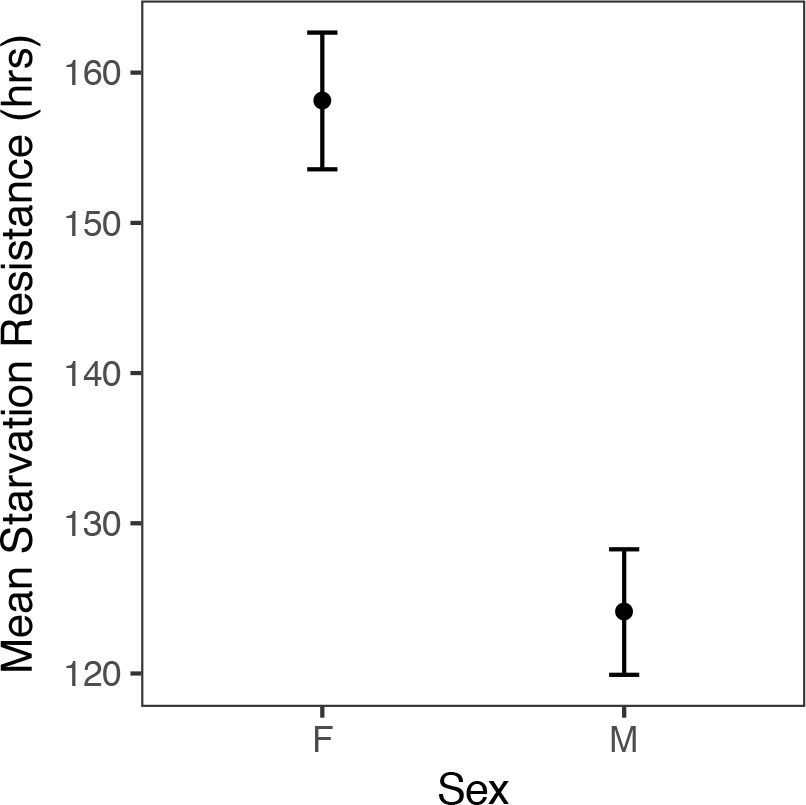
Mean starvation resistance (±95% CI) for males and females in the DGRP (F_1,334_ = 118.21, p < 0.0001).

**Figure S5.**
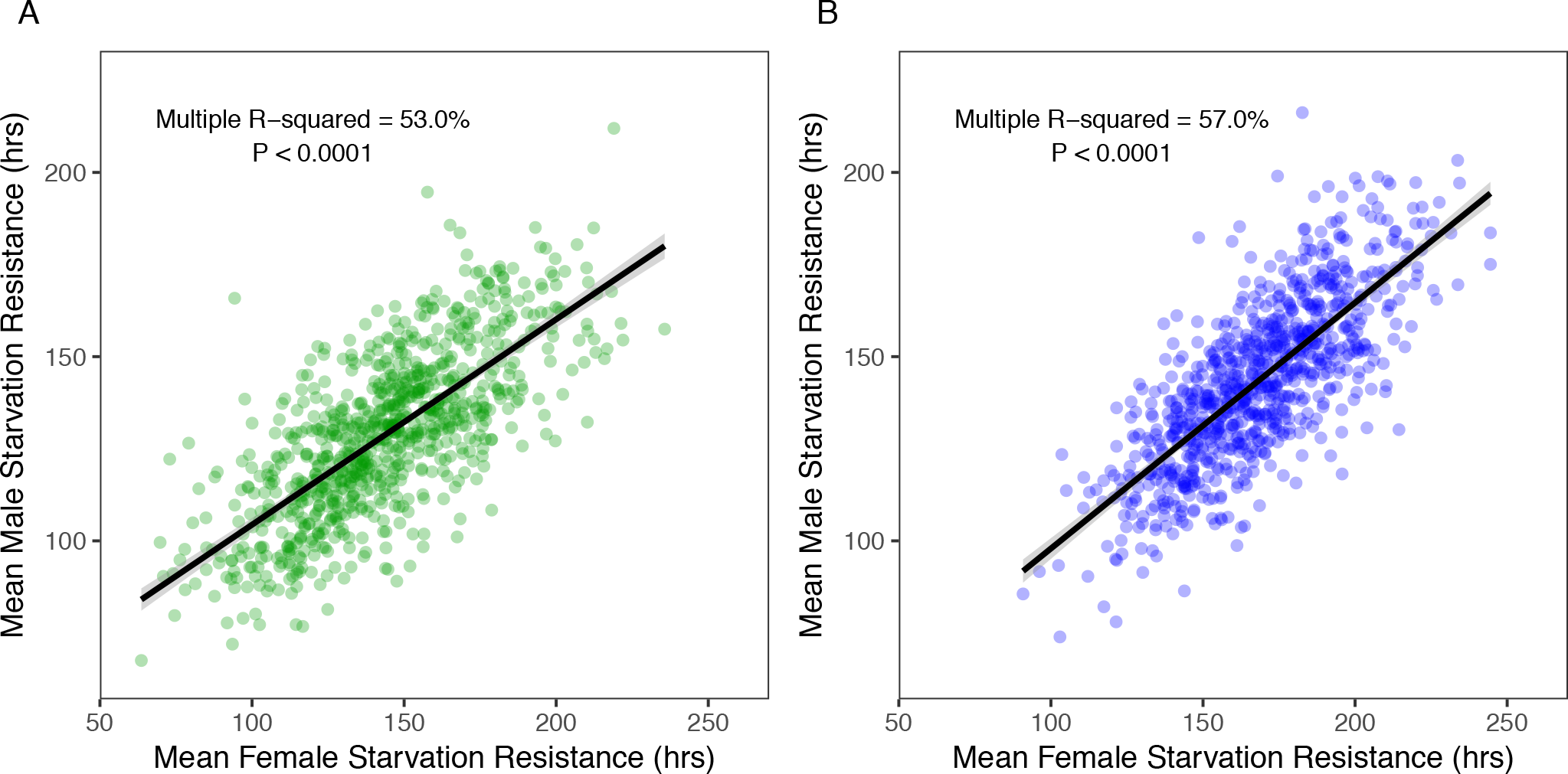
Correlation between sex-specific mean starvation resistance for the DSPR pA mapping panel (A) and pB mapping panel (B). Sex-specific responses were significantly correlated for both panels (pA: F_1,859_ = 968.5, P < 0.0001; pB: F_1,860_ = 1138, p < 0.0001). Grey shading around the regression line in both plots indicates the 95% confidence interval.

**Figure S6.**
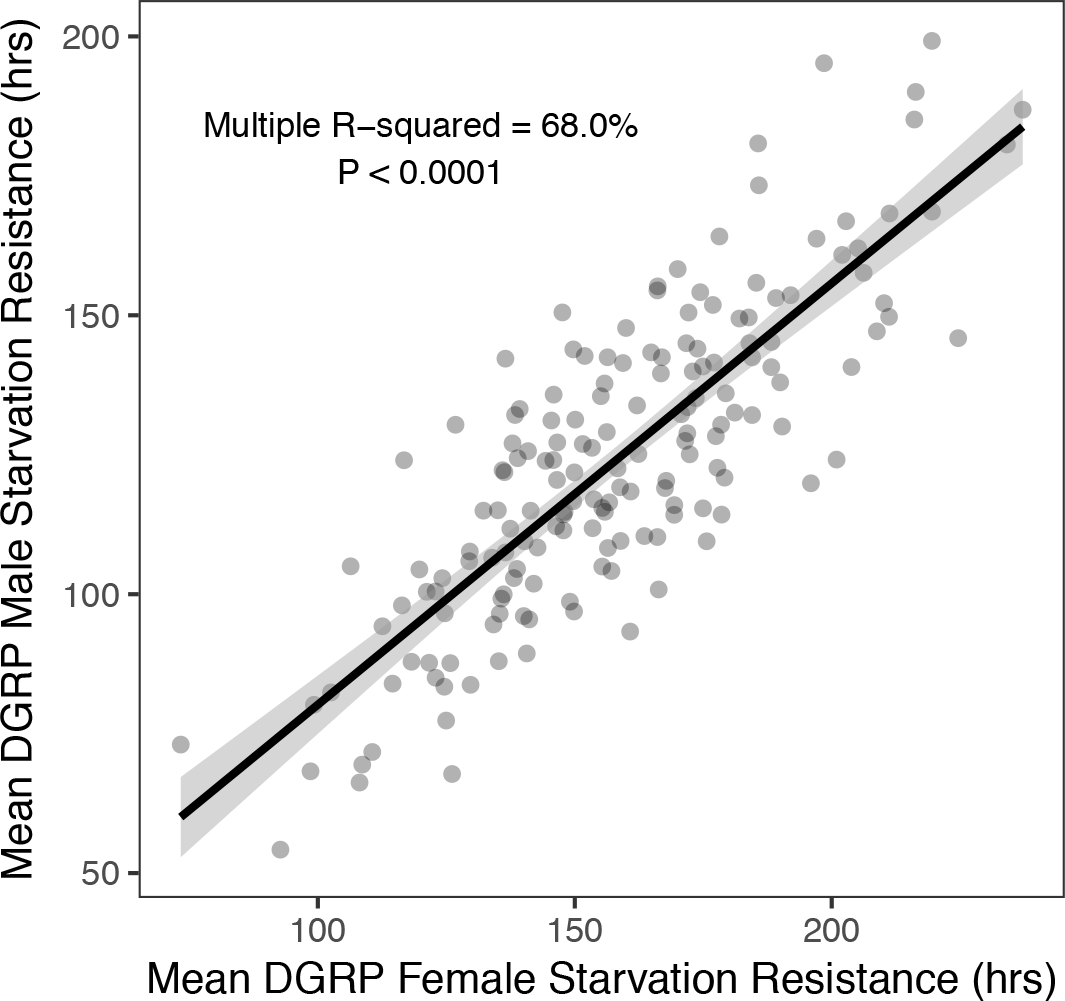
Male and female mean starvation resistance was significantly correlated in the DGRP (F_1,334_ = 118.21, P < 0.0001). Grey shading around the regression line indicates the 95% confidence interval.

**Figure S7.**
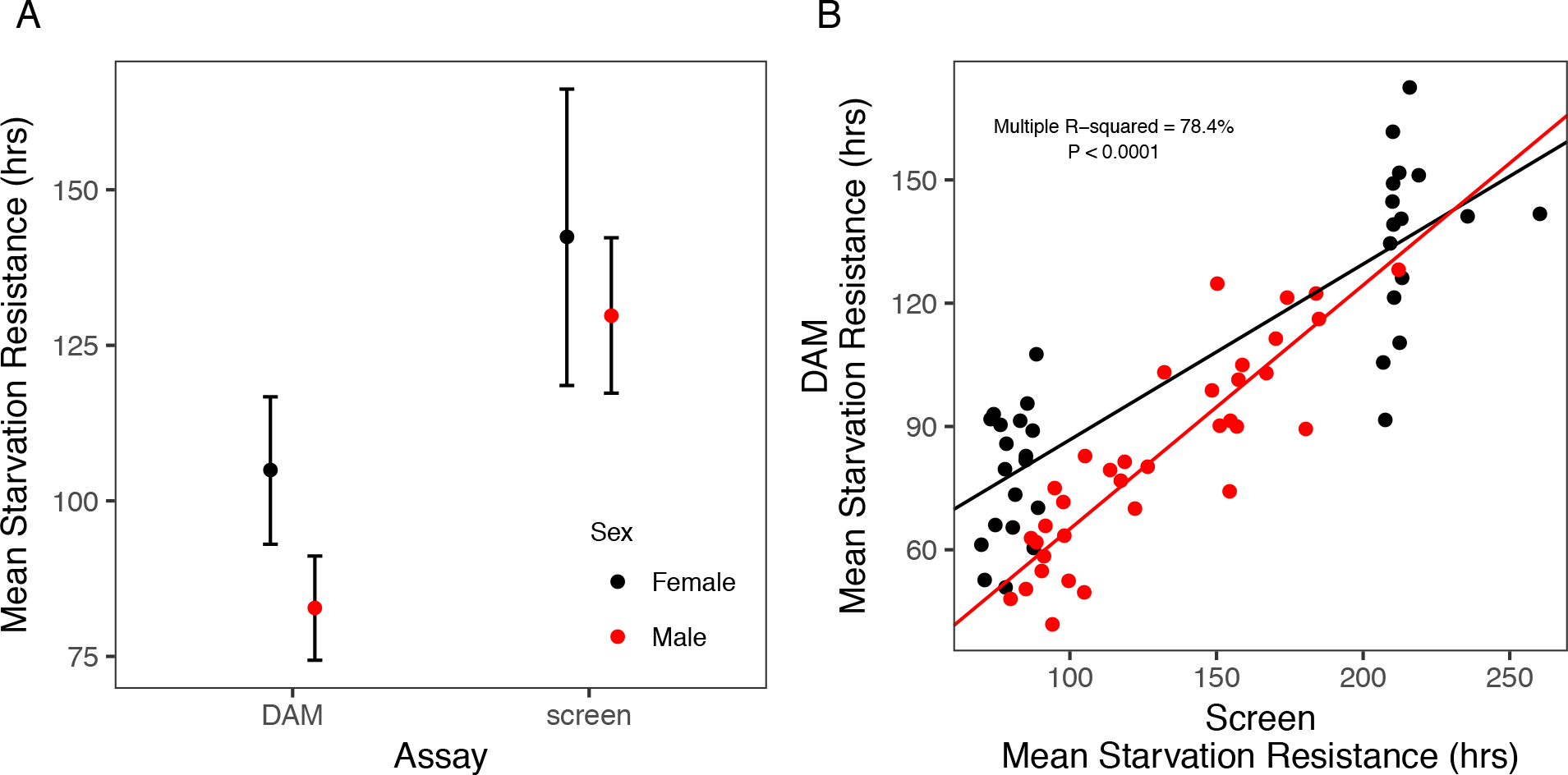
A. Starvation resistance was significantly higher in the large-scale starvation screen of all DSPR RILs compared to the DAM (Drosophila Activity Monitor) starvation assay for the selected subset of RILs (Assay: F_1,136_ = 31.60, p < 0.0001). Mean starvation resistance across RIL means is presented (±95% CI). B. Mean starvation resistance measured in the large-scale starvation resistance screen (x-axis) was correlated with mean starvation resistance measured in the DAM assay (y-axis) in the DSPR (Females: β = 0.43 ± 0.04, t = 9.7, p < 0.0001, R^2 =^ 73.9%; Males: β = 0.59 ± 0.05, t = 10.9, p < 0.0001, R^2 =^ 78.3%). The multiple R^2^ value in the plot includes the interaction between starvation resistance measured under different assay conditions with sex.

**Figure S8.**
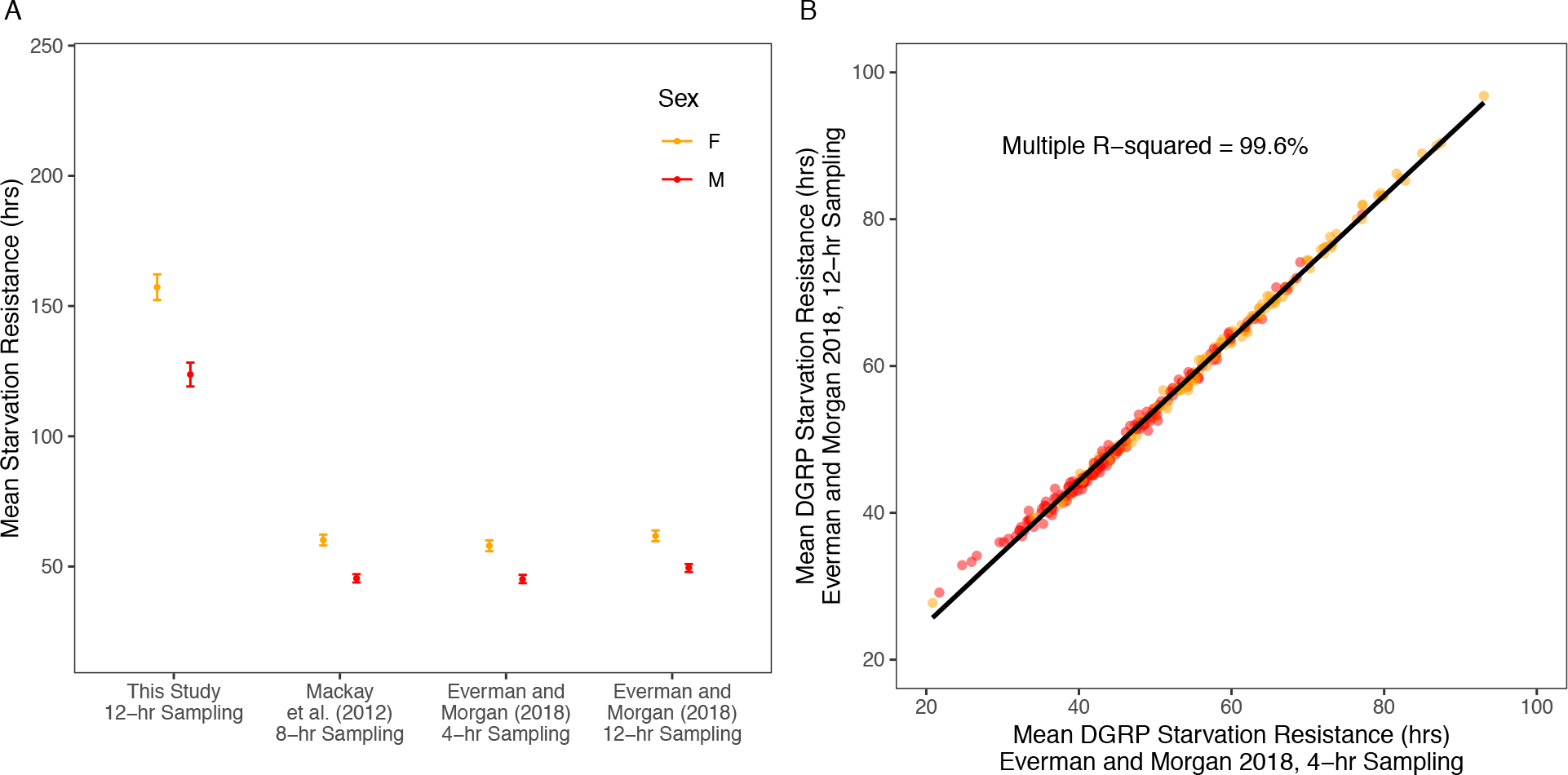
A. Mean starvation resistance (±95% CI) was significantly higher in this study compared to Mackay et al. (2012) and Everman and Morgan (2018) (F_2.532_ = 1457.5, P < 0.0001). The increased mean and variation in starvation resistance observed in this study was not driven by differences in the frequency at which survival was assessed, since a re-analysis of data from Everman and Morgan (2018) with a longer interval between fly counting events matching the interval from the present study, revealed essentially no difference in the phenotypes assayed. B. Mean starvation resistance by line and sex measured according to the 4-hr sampling interval was highly correlated to our re-analysis of the Everman and Morgan (2018) data using a 12-hr sampling interval.

**Figure S9.**
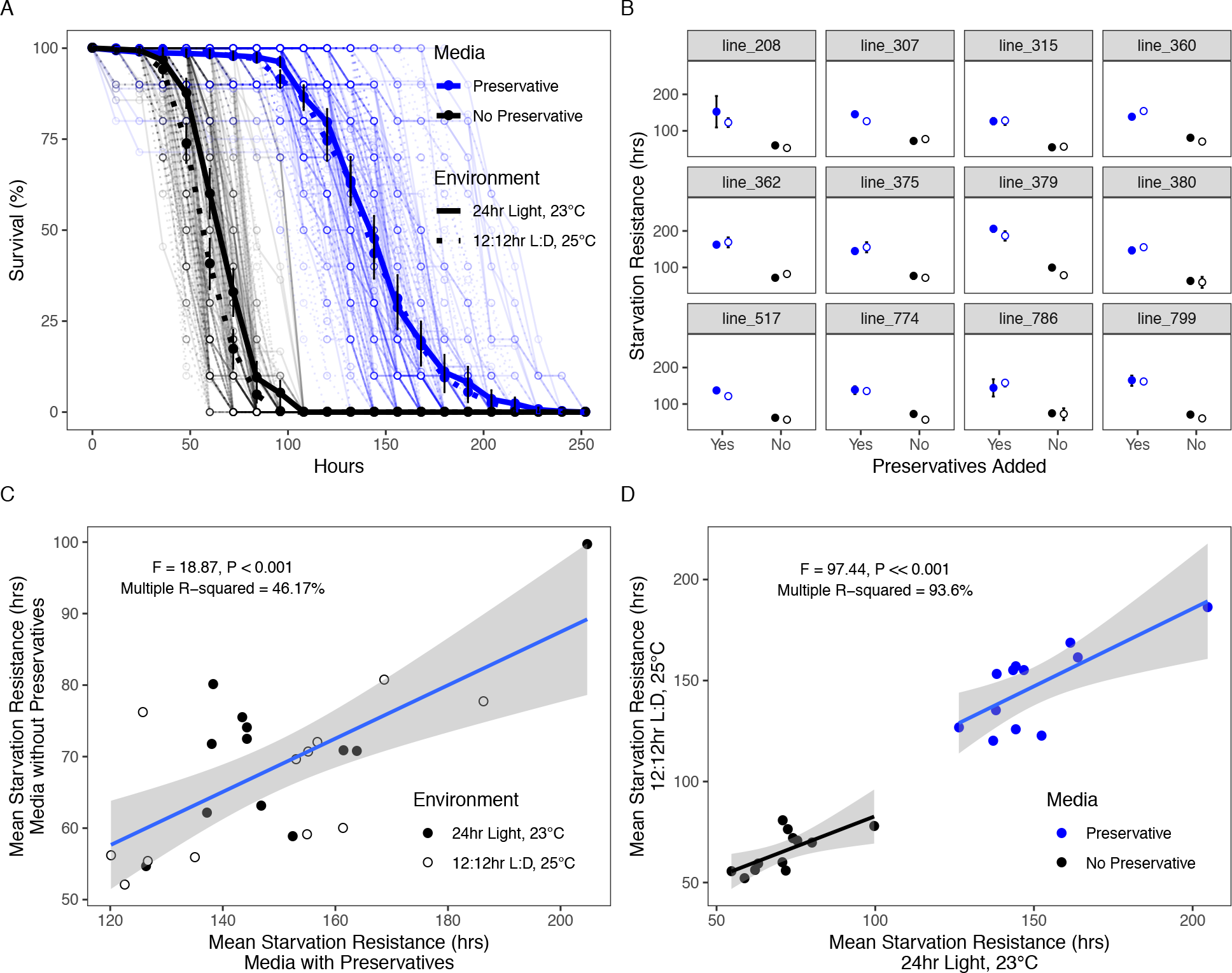
Flies maintained on starvation media with preservatives lived much longer than flies on starvation media without preservatives, regardless of environmental conditions (24-hr light, 23°C as used in the large-scale starvation screen vs. 12:12hr L:D, 25°C as used in Mackay et al. (2012) and Everman and Morgan (2018)). This was tested in 12 randomly-selected DGRP lines. A. Percent survival per vial was different between the two media treatments, and differed slightly due to environment, but only when media did not contain preservatives. Black lines and points indicate media with no preservatives; blue lines and points indicate media with preservatives; solid lines indicate the 24hr Light, 23°C environment; dashed lines indicate the 12:12hr L:D, 25°C treatment. The bold points and lines for each treatment indicate the overall mean (±95% CI) survival of each treatment group at each 12-hr assessment point. B. Mean (±95% CI) starvation resistance for each of the 12 randomly selected DGRP lines was rarely influenced by the environment treatment (closed symbols = 24hr Light, 23°C; open symbols = 12:12hr L:D, 25°C), but media preservatives consistently resulted in higher survival for each DGRP line. C. Mean starvation resistance by DGRP line was significantly correlated between the two media treatments. D. DGRP line, preservatives in the media, and environmental conditions together explained nearly all phenotypic variation in starvation resistance. The full reporting of variance components is presented in Table S6. In C and D, grey shading represents the 95% CI of the regression.

**Figure S10.**
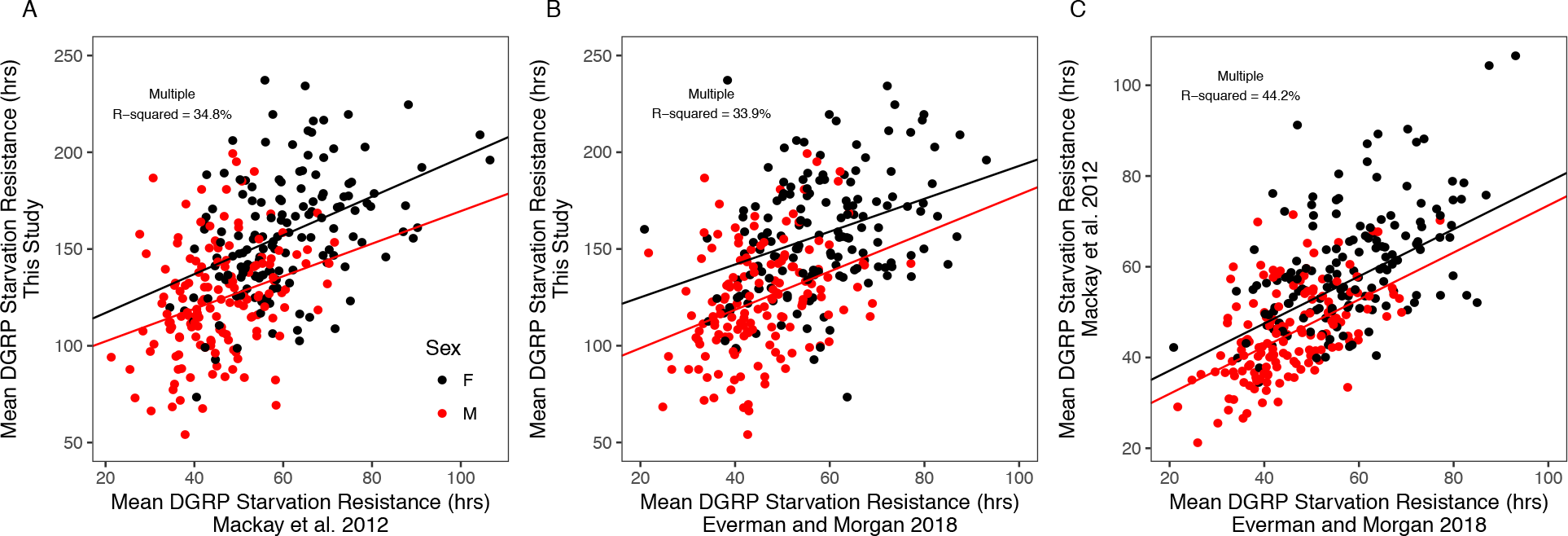
Correlation between sex-specific mean starvation resistance in the DGRP panel for 150 lines that overlap between this study, Mackay et al. 2012, and Everman and Morgan 2018. Red points indicate males and black points indicate females. All comparisons showed that the three independent measures of starvation resistance were significantly correlated (A: F_3,296_ = 52.69, p < 0.0001; B: F_3, 296_ = 50.66, p < 0.0001; C: F_3.296_ = 78.18, p < 0.0001).

**Figure S11.**
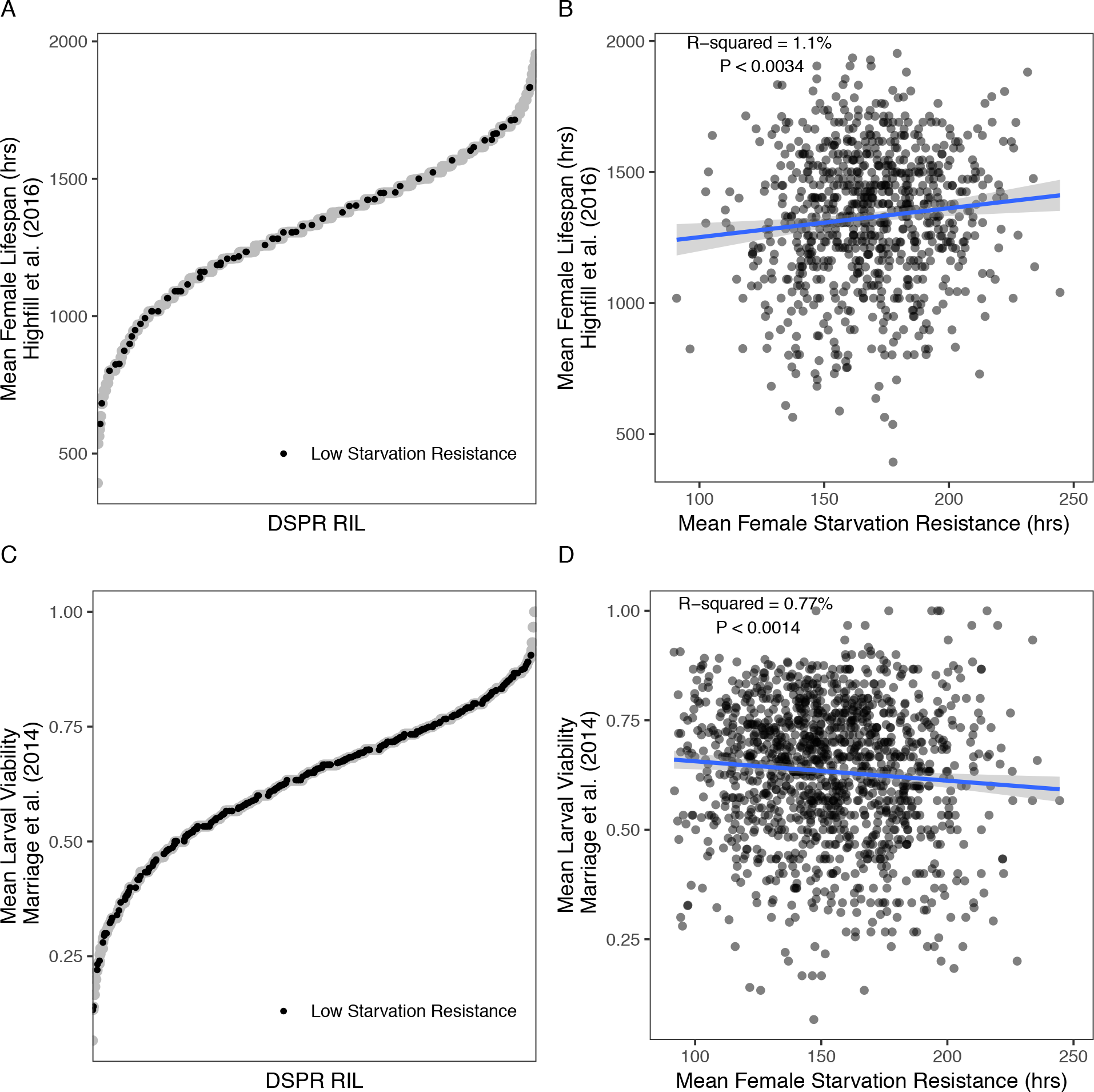
DSPR RILs with low starvation resistance ranged from low to high for other measures of fitness. A. Distribution of mean female lifespan in DSPR pB RILs (Highfill et al. (2016). RILs with low female starvation resistance (in the bottom 25% of the distribution) are shown in solid black symbols, while other RILs are shown in gray. B. The correlation between mean female starvation resistance and mean female lifespan is minimal, suggesting there is no association between low starvation resistance and reduced lifespan. C. Distribution of mean larval viability measured as the proportion of 1^st^ instar larvae reared under control conditions that emerged as adults (Marriage et al. (2014). RILs with low female starvation resistance (in the bottom 25% of the distribution) are shown in solid black symbols, while other RILs are shown in gray. D. The weak correlation between mean female starvation resistance and mean larval viability again suggests that there is no association between low starvation resistance and low larval viability. We also fail to find strong associations between starvation resistance and other measures of fitness in the DGRP (Fig S12).

**Figure S12.**
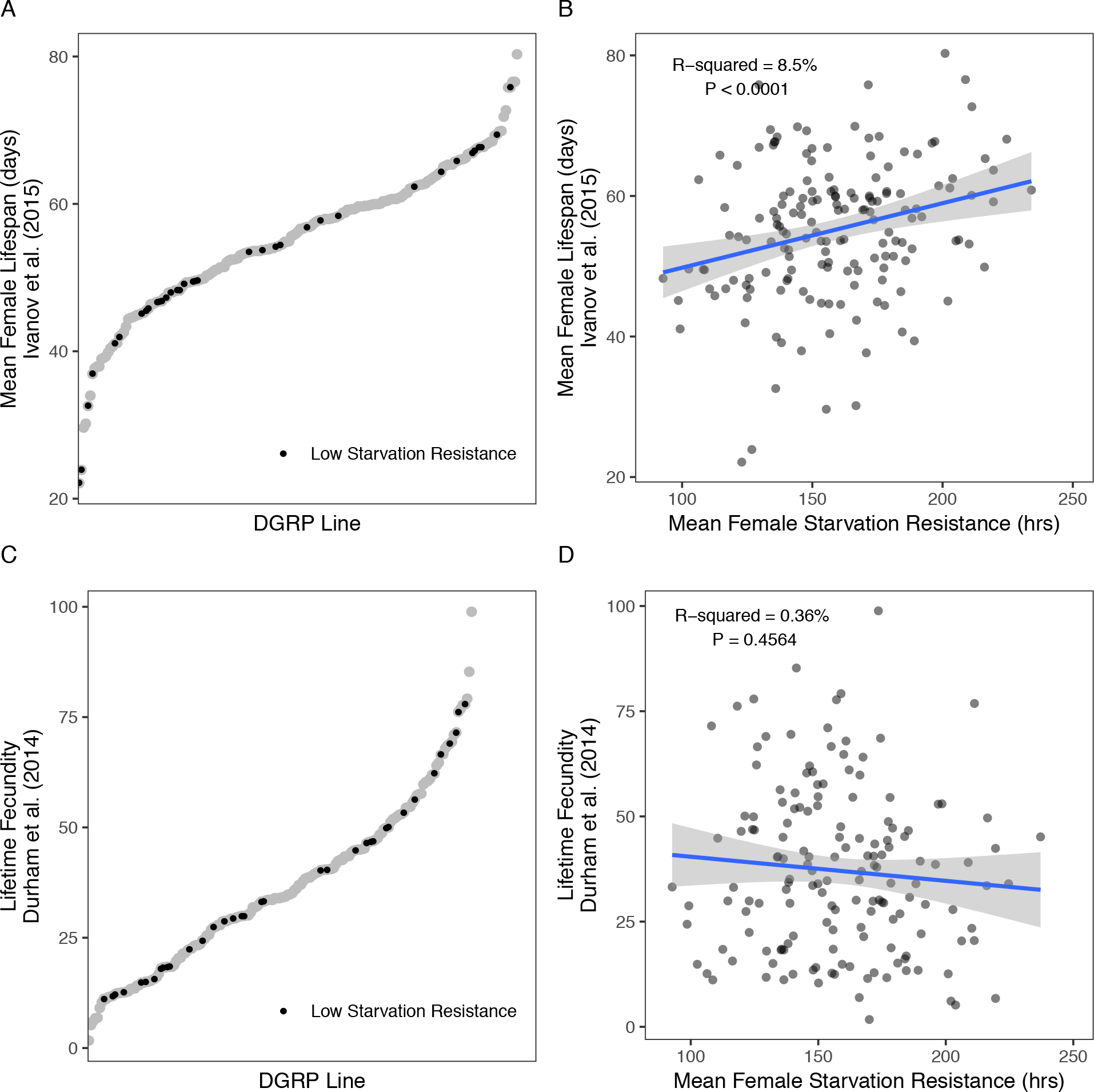
DGRP Lines with low starvation resistance ranged from low to high for other measures of fitness. A. Distribution of mean female lifespan in the DGRP (Ivanov et al. (2015). Lines with low female starvation resistance (in the bottom 25% of the distribution) are shown in solid black symbols, while other lines are shown in gray. B. The correlation between mean female starvation resistance and mean female lifespan is minimal, suggesting there is little association between low starvation resistance and reduced lifespan. C. Distribution of lifetime fecundity (Durham et al. (2014). Lines with low female starvation resistance (in the bottom 25% of the distribution) are shown in solid black symbols, while other lines are shown in gray. D. The weak correlation between mean female starvation resistance and fecundity suggests that there is no association between low starvation resistance and this measure of fitness.

**Figure S13.**
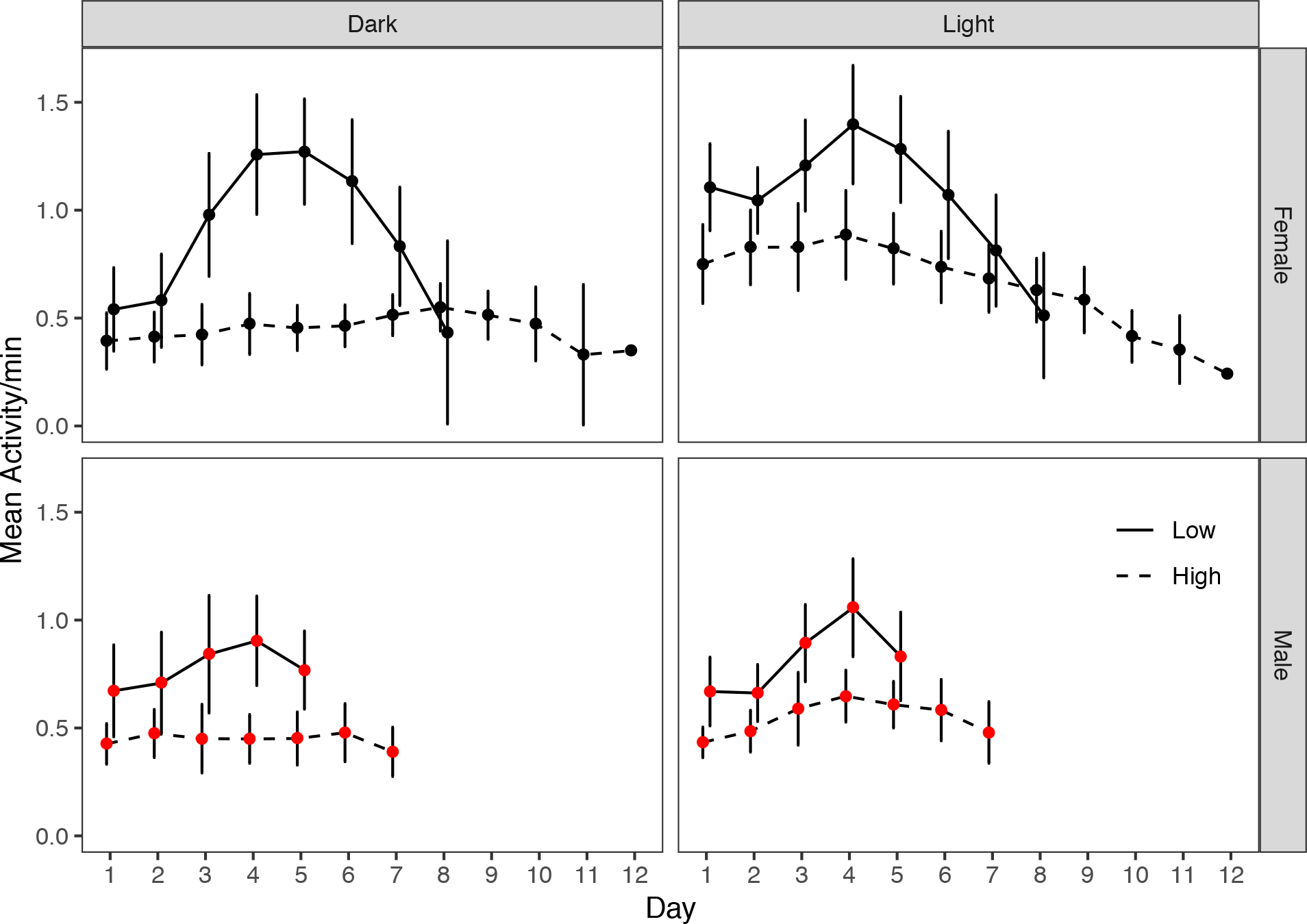
Mean activity levels for females (top panel, black points) and males (bottom panel, red points) (±95% CI) across the 12-hr daily light or dark periods during starvation in the DAM (Drosophila Activity Monitor) assay until death. Low starvation resistance RILs (solid line) tended to be more active during starvation compared to high starvation resistance RILs (dashed line) under both light and dark conditions. Light status (dark versus light) influenced the overall activity level of females but did not influence male activity. Data were analyzed with a repeated measures ANOVA; results are presented in Table S8. Similar to the pre-starvation period (Fig 4), waking activity levels of individuals during the DAM starvation experiment were primarily driven by starvation resistance rank in both sexes (females: F_1,8_ = 14.87, p < 0.01; males: F_1,6_ = 13.87, p < 0.01; Table S8). Light status did not influence activity between days for either sex (females: F_1,8_ = 0.84, p = 0.39; males: F_1,6_ = 0.23, p = 0.65; Table S8). However, light status did significantly influence activity in females within each day (females: F_1,8_ = 43.09, p < 0.0001; Table S8), indicating that female activity in both high and low starvation resistance RILs was consistently higher during times when lights were on. Male activity of high and low starvation resistance RILs tended to remain constant under different light conditions. The larger influence of light status on females is consistent with patterns observed in activity during the pre-starvation period (Fig 4).

**Figure S14.**
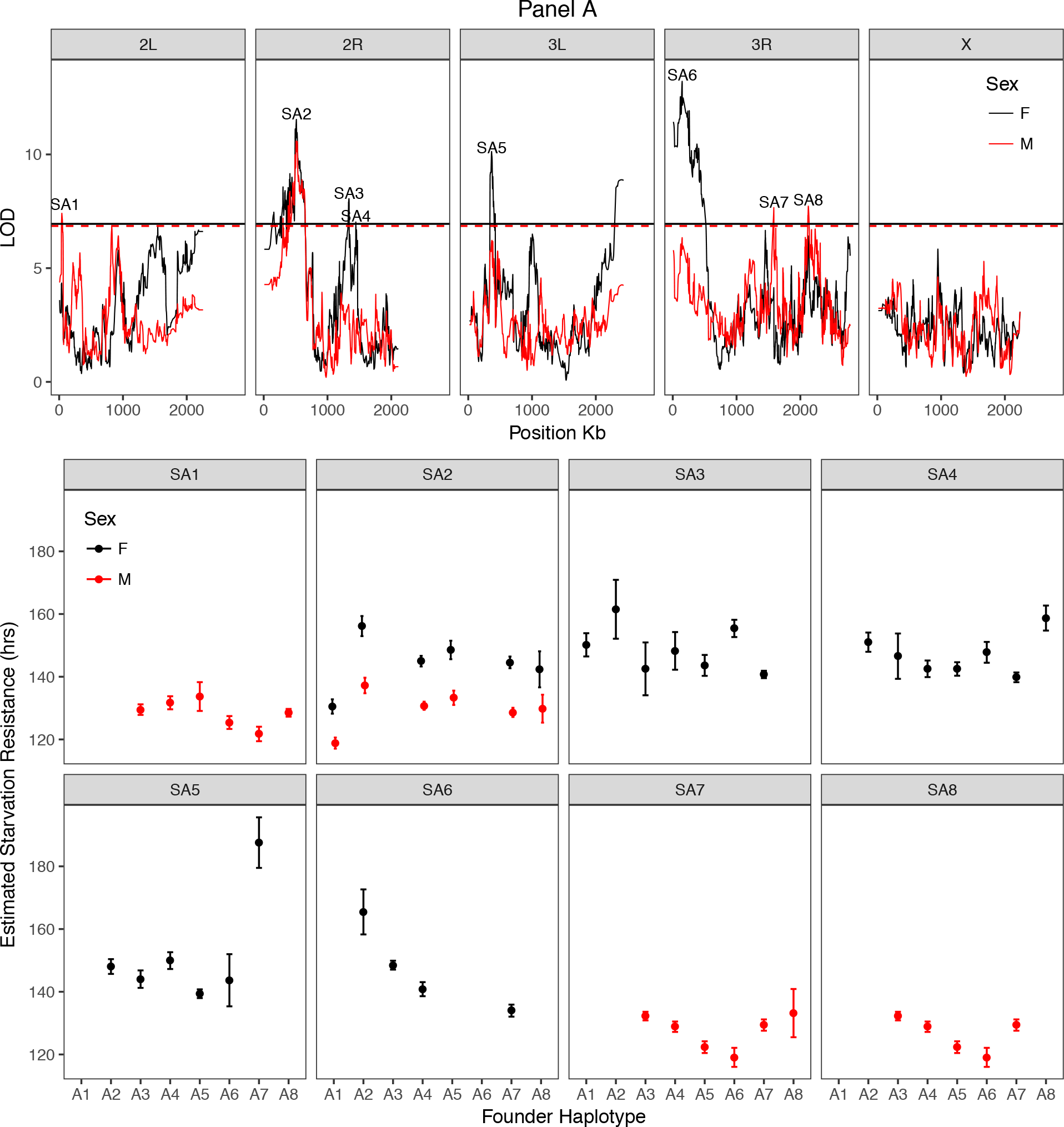
Starvation resistance QTL and estimated allele effects at each QTL. Data are presented as RIL means (±SE) for estimated starvation resistance when the founder haplotype was present in more than 5 RILs. As has been seen in a number of studies using the DSPR and other multiparental populations (King et al. 2012b; Giraud et al. 2014; Najarro et al. 2015), the estimated phenotypic effects of each founder haplotype suggest that multiple alleles may be present at our starvation QTL, since the effects do not fall into two clear “high” and “low” classes.

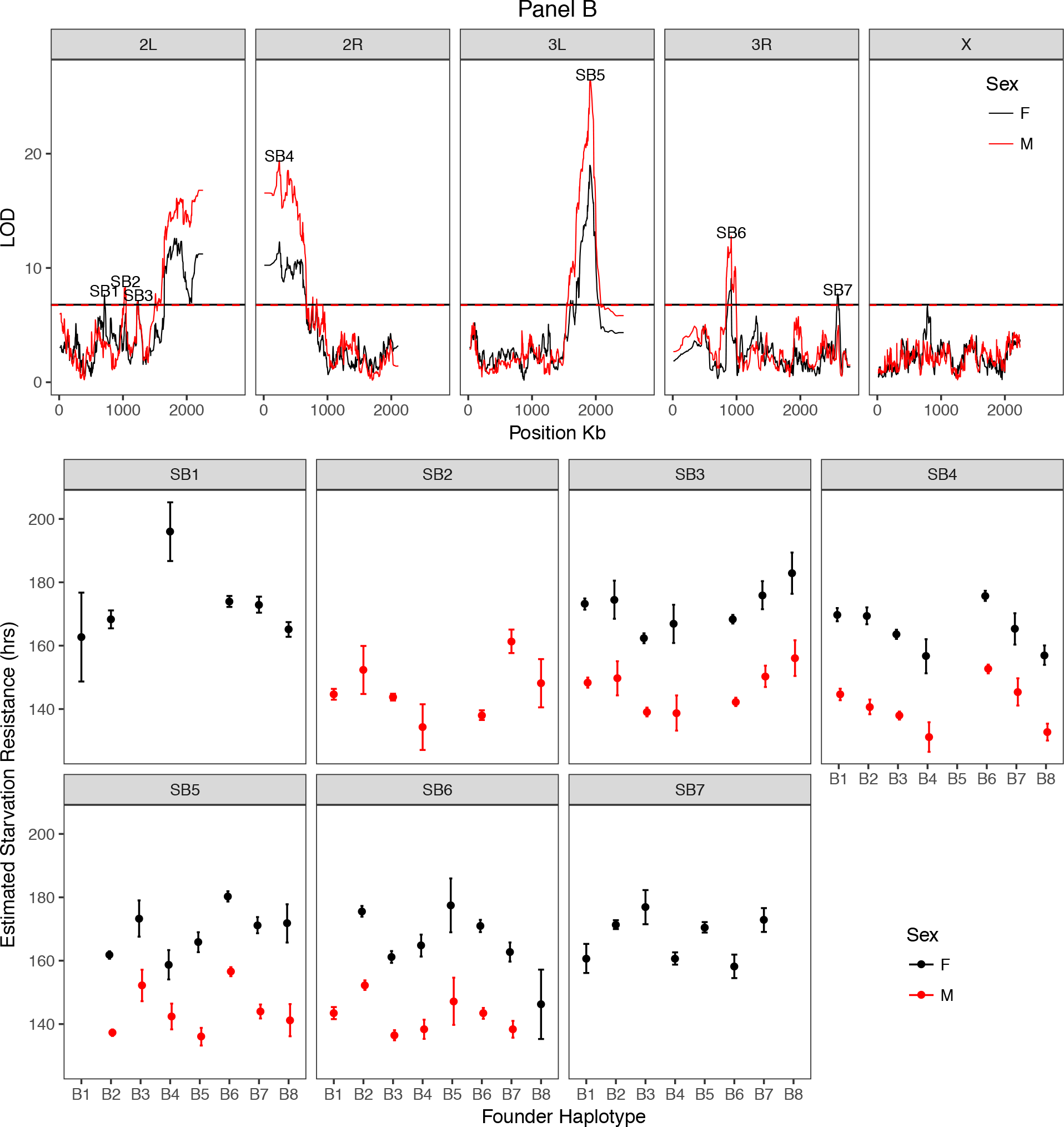

**Figure S15.**
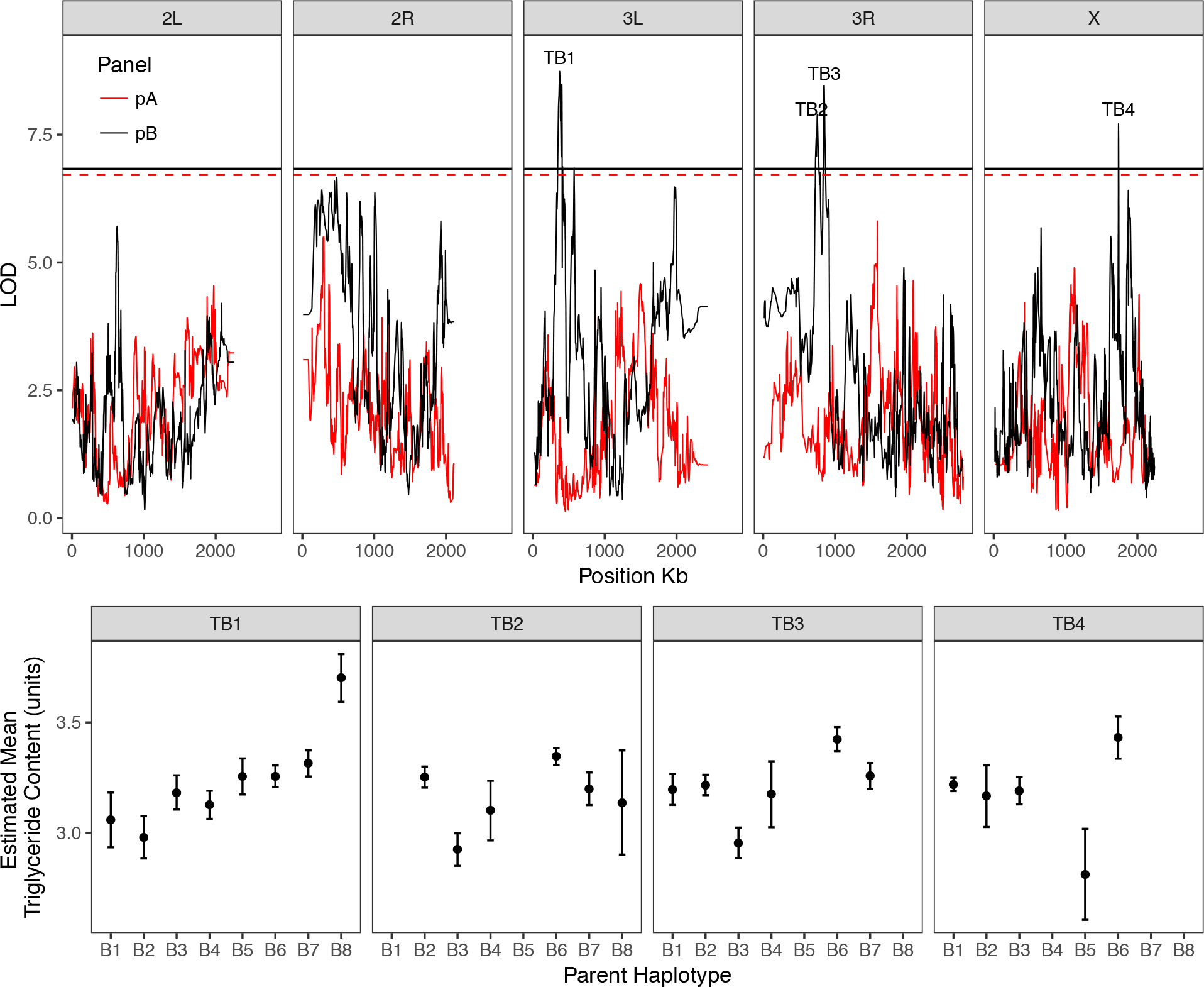
Triglyceride level QTL and estimated allele effects of founders at each QTL. Data are presented as RIL means (±SE) for estimated triglyceride level when the founder haplotype was present in more than 5 RILs. Similar to starvation resistance, the estimated phenotypic effects of each founder haplotype suggest that multiple alleles may be present at our triglyceride QTL.

**Figure S16.**
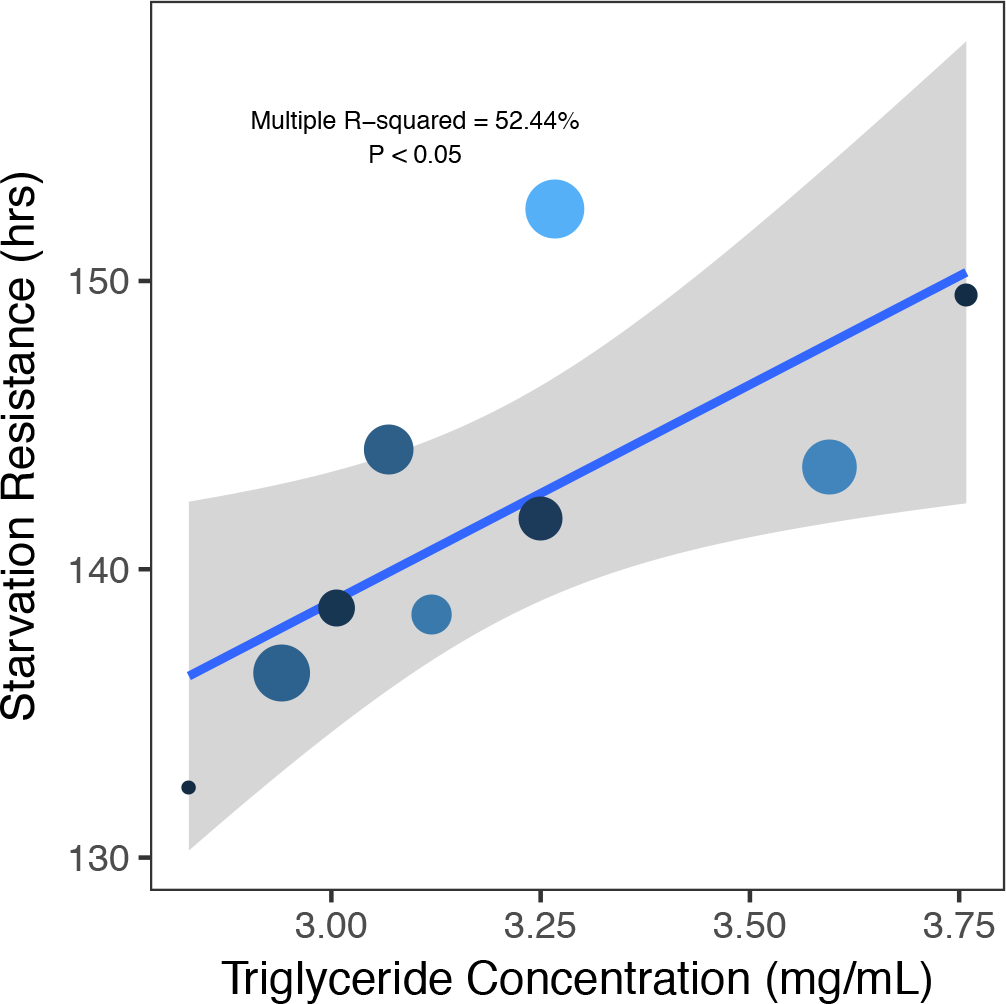
Triglyceride level and starvation resistance were correlated after accounting for variation due to founder haplotype at the overlapping peaks TB3 and SB6 (F_1,7_ = 7.72, P < 0.05). Data presented are averages for each founder haplotype in the pB panel, including “NA” for RILs that could not be assigned with confidence to a known haplotype. Point size relates the number of RILs per haplotype for the starvation resistance peak (smallest = 1 RIL; largest = 193 RILs); point color relates the number of RILs per haplotype for the triglyceride level peak (black = 1 RIL; lightest blue = 181 RILs). Grey shading around the regression line indicates the 95% confidence interval.

**Figure S17.**
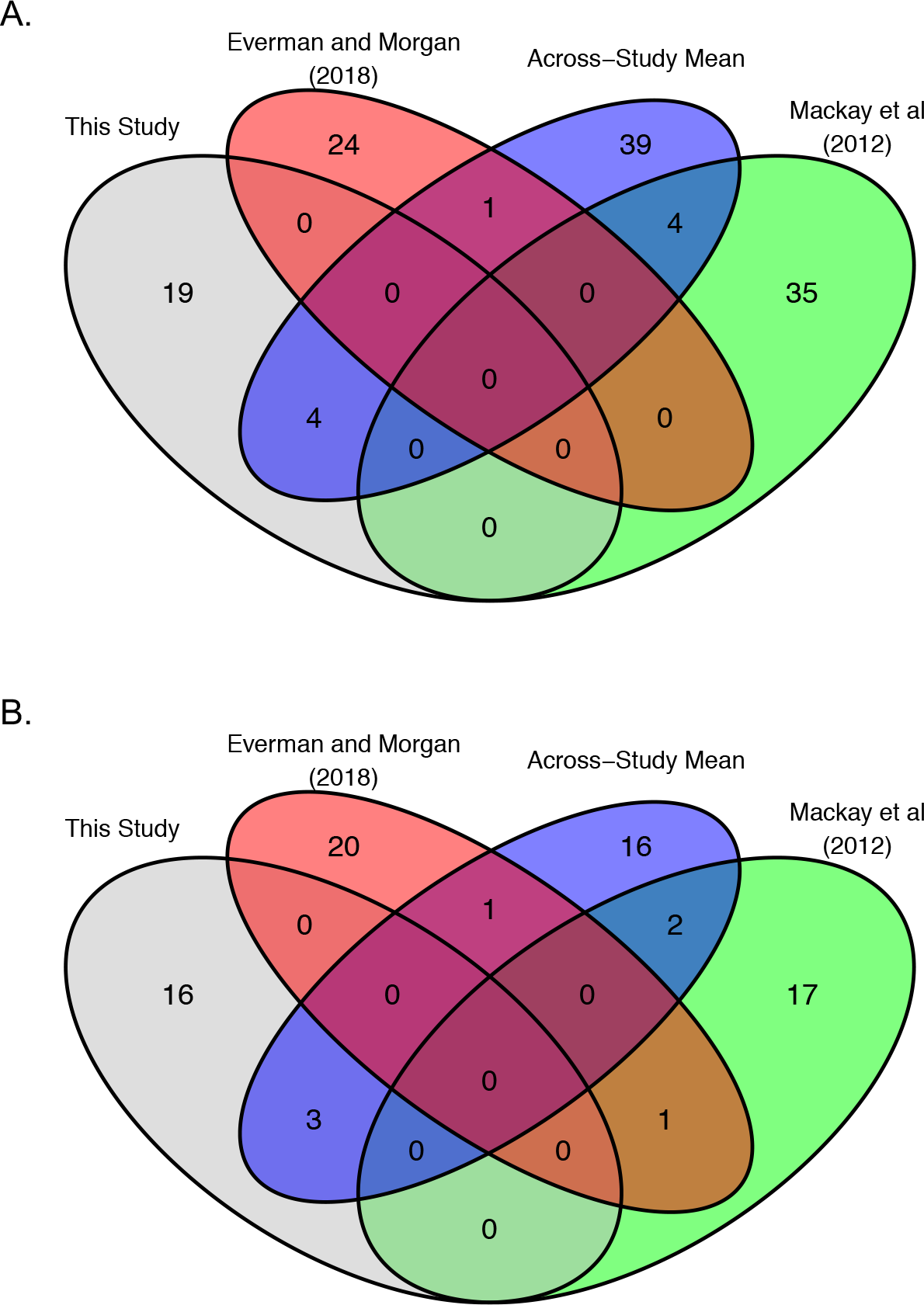
Overlap in SNPs associated with starvation resistance for each DGRP dataset using the P < 10^− 5^ significance threshold. Overlap between data sets was minimal. Plot A presents results for females; plot B presents results for males.

**Figure S18.**
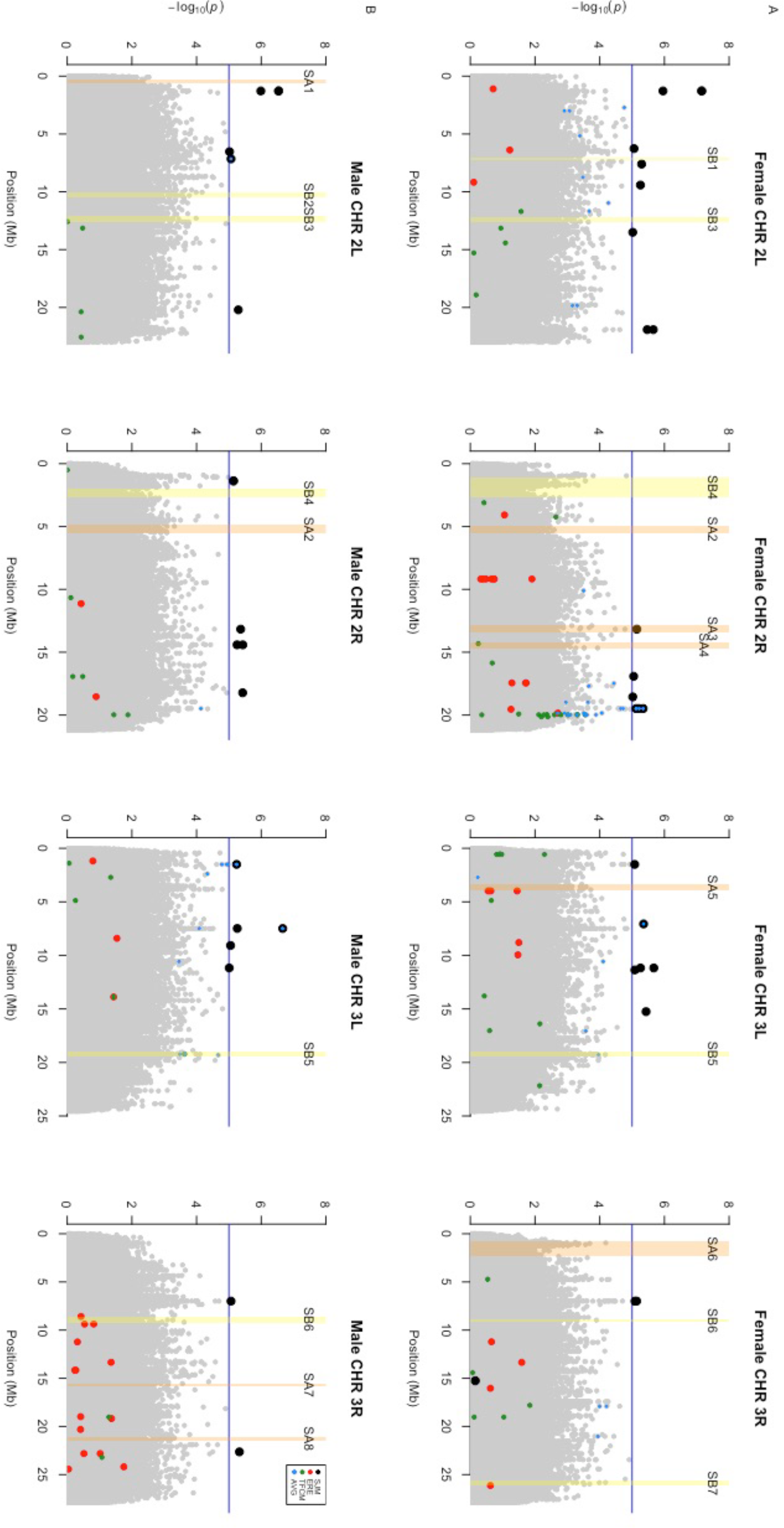
Manhattan plots of mean starvation resistance in the DGRP with SNPs that were associated with starvation resistance in previous studies and intervals of sex-specific QTL identified for starvation resistance in the DSPR highlighted. Plots are broken up by chromosome arm in A for females and in B for males. In all plots, points highlighted in black indicate SNPs that are associated with starvation resistance in the DGRP from data obtained in this study; red points indicate SNPs associated with starvation resistance in Everman and Morgan 2018; green points indicate SNPs associated with starvation resistance in Mackay et al. 2012; blue points indicate SNPs associated with starvation resistance averaged across the three datasets. A genomewide significance threshold of P < 10^−5^ is shown with the blue line. Yellow shaded boxes and labels correspond to QTL intervals around peaks mapped in the pB DSPR panel; orange shaded boxes correspond to QTL intervals around peaks mapped in the pA DSPR panel.

**Figure S18.**
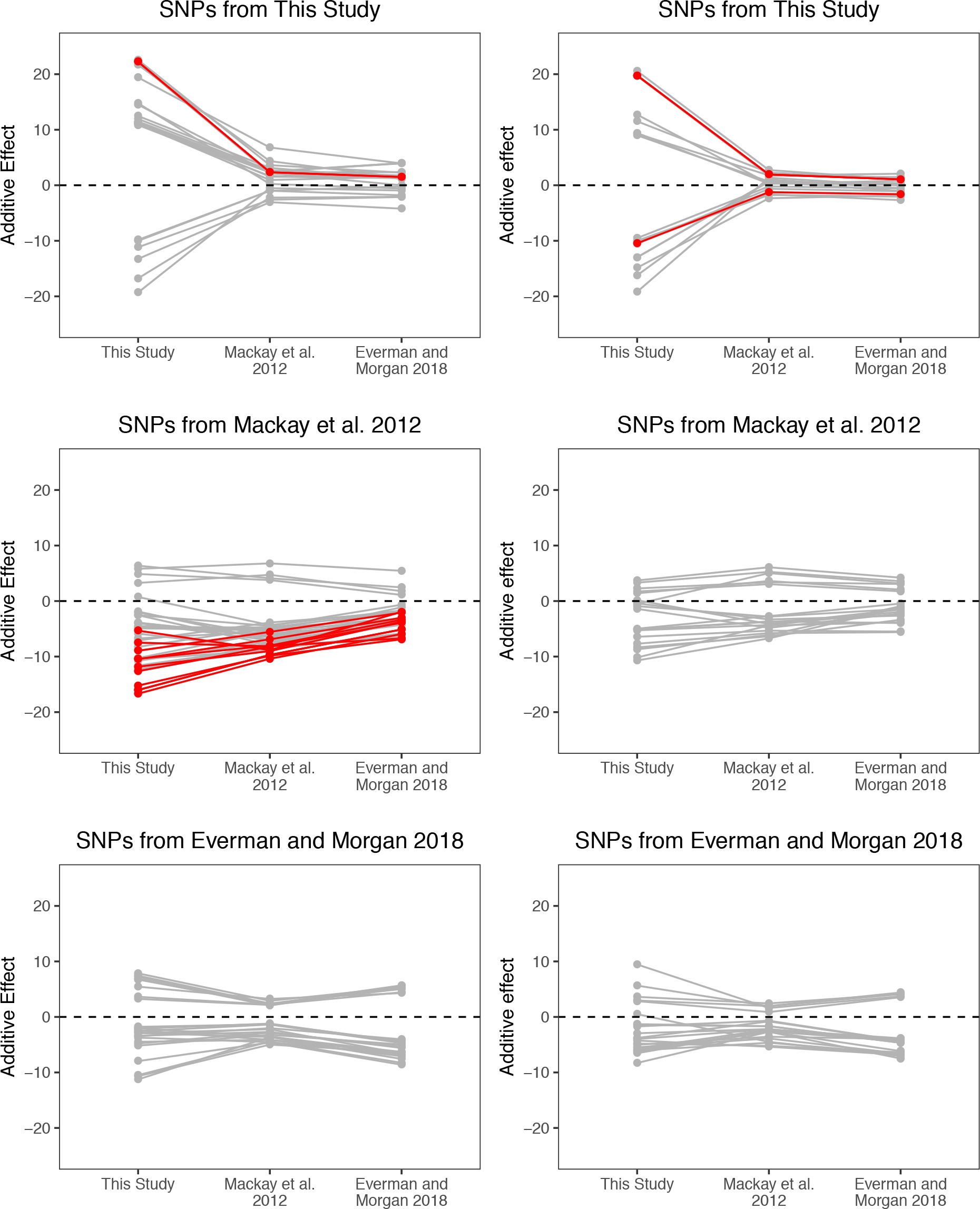
Additive effects of SNPs associated with starvation resistance (at P < 10^−5^) in each study, along with their additive effects estimated in the other two studies. Female data is presented in the left column of plots; male data is presented in the right column of plots. SNPs that passed the FDR threshold of 0.2 are highlighted in red. Generally, SNPs had similar effects (of the same +/-sign) on starvation resistance in all three experiments.

## Supplemental Tables

**Table S1.** Analysis of variance of the effect of DSPR rearing density on starvation resistance.

| Source | df | SS | MS | F | P | % Var. Exp. |
| --- | --- | --- | --- | --- | --- | --- |
| Density | 1 | 1758.00 | 1758.20 | 18.15 | < 0.0001 | 0.90 |
| DSPR RIL | 19 | 148747 | 7828.80 | 80.83 | << 0.0001 | 80.2 |
| Density x DSPR RIL | 19 | 8905 | 468.70 | 4.84 | < 0.0001 | 4.80 |
| Residual | 269 | 26054 | 96.90 |  |  |  |

**Table S2.** Analysis of variance of mean starvation resistance in the DSPR.

| Source | df | SS | MS | F | P |
| --- | --- | --- | --- | --- | --- |
| Subpopulation<sup>a</sup> | 3 | 330782 | 110261 | 183.702 | < 0.0001 |
| Sex | 1 | 342197 | 342197 | 570.121 | < 0.0001 |
| Subpopulation x Sex | 3 | 32982 | 10994 | 18.317 | < 0.0001 |
| Residual | 3440 | 2064747 | 600 |  |  |

<sup>a</sup> We note that since most RILs from a given subpopulation were tested in the same batch, batch effects may contribute to some of the subpopulation-to-subpopulation differences we report.

**Table S3.** Analysis of variance of mean triglyceride level in the DSPR.

| **Source** | **df** | **SS** | **MS** | **F** | **P** |
| --- | --- | --- | --- | --- | --- |
| Subpopulation | 3 | 46.81 | 15.6020 | 37.099 | < 0.0001 |
| Residual | 935 | 393.22 | 0.4206 |  |  |

**Table S4.** Stratification of the top 50 SNPs associated with starvation resistance in the DGRP across five frequency bins.

| Study | Sex | Allele Frequency Bin | No. SNPs |
| --- | --- | --- | --- |
| This Study | F | 0.05 - 0.1 | 14 |
| This Study | F | > 0.1 - 0.2 | 9 |
| This Study | F | > 0.2 - 0.3 | 13 |
| This Study | F | > 0.3 - 0.4 | 6 |
| This Study | F | > 0.4 - 0.5 | 8 |
| This Study | M | 0.05 - 0.1 | 11 |
| This Study | M | > 0.1 - 0.2 | 12 |
| This Study | M | > 0.2 - 0.3 | 13 |
| This Study | M | > 0.3 - 0.4 | 3 |
| This Study | M | > 0.4 - 0.5 | 11 |
| Mackay *et al.* 2012 | F | 0.05 - 0.1 | 28 |
| Mackay *et al.* 2012 | F | > 0.1 - 0.2 | 11 |
| Mackay *et al.* 2012 | F | > 0.2 - 0.3 | 5 |
| Mackay *et al.* 2012 | F | > 0.3 - 0.4 | 2 |
| Mackay *et al.* 2012 | F | > 0.4 - 0.5 | 4 |
| Mackay *et al.* 2012 | M | 0.05 - 0.1 | 21 |
| Mackay *et al.* 2012 | M | > 0.1 - 0.2 | 13 |
| Mackay *et al.* 2012 | M | > 0.2 - 0.3 | 7 |
| Mackay *et al.* 2012 | M | > 0.3 - 0.4 | 2 |
| Mackay *et al.* 2012 | M | > 0.4 - 0.5 | 7 |
| Everman and Morgan 2018 | F | 0.05 - 0.1 | 11 |
| Everman and Morgan 2018 | F | > 0.1 - 0.2 | 13 |
| Everman and Morgan 2018 | F | > 0.2 - 0.3 | 10 |
| Everman and Morgan 2018 | F | > 0.3 - 0.4 | 8 |
| Everman and Morgan 2018 | F | > 0.4 - 0.5 | 8 |
| Everman and Morgan 2018 | M | 0.05 - 0.1 | 18 |
| Everman and Morgan 2018 | M | > 0.1 - 0.2 | 13 |
| Everman and Morgan 2018 | M | > 0.2 - 0.3 | 12 |
| Everman and Morgan 2018 | M | > 0.3 - 0.4 | 4 |
| Everman and Morgan 2018 | M | > 0.4 - 0.5 | 3 |

**Table S5.** Analysis of variance of mean starvation resistance in the DGRP.

| Source | df | SS | MS | F | P |
| --- | --- | --- | --- | --- | --- |
| Sex | 1 | 97213 | 97213 | 118.21 | < 0.0001 |
| Residual | 334 | 274680 | 822 |  |  |

**Table S6.** Analysis of variance of the effect of preservatives and environment on starvation resistance in the DGRP.

| Source | df | SS | MS | F value | P | % Var. Exp. |
| --- | --- | --- | --- | --- | --- | --- |
| Environment | 1 | 2400 | 2400 | 16.71 | < 0.0001 | 0.30 |
| Preservatives | 1 | 604184 | 604184 | 4205.70 | < 0.0001 | 81.17 |
| DGRP Line | 11 | 63653 | 5787 | 40.28 | < 0.0001 | 8.55 |
| Environment x Preservatives | 1 | 155 | 155 | 1.08 | 0.30 | 0.02 |
| Environment x DGRP Line | 11 | 7289 | 663 | 4.61 | < 0.0001 | 0.98 |
| Preservatives x DGRP Line | 11 | 14718 | 1338 | 9.31 | < 0.0001 | 1.98 |
| Environment x Preservatives x DGRP Line | 11 | 4961 | 451 | 3.14 | < 0.001 | 0.67 |
| Residual | 327 | 46976 | 144 |  |  |  |

**Table S7.**
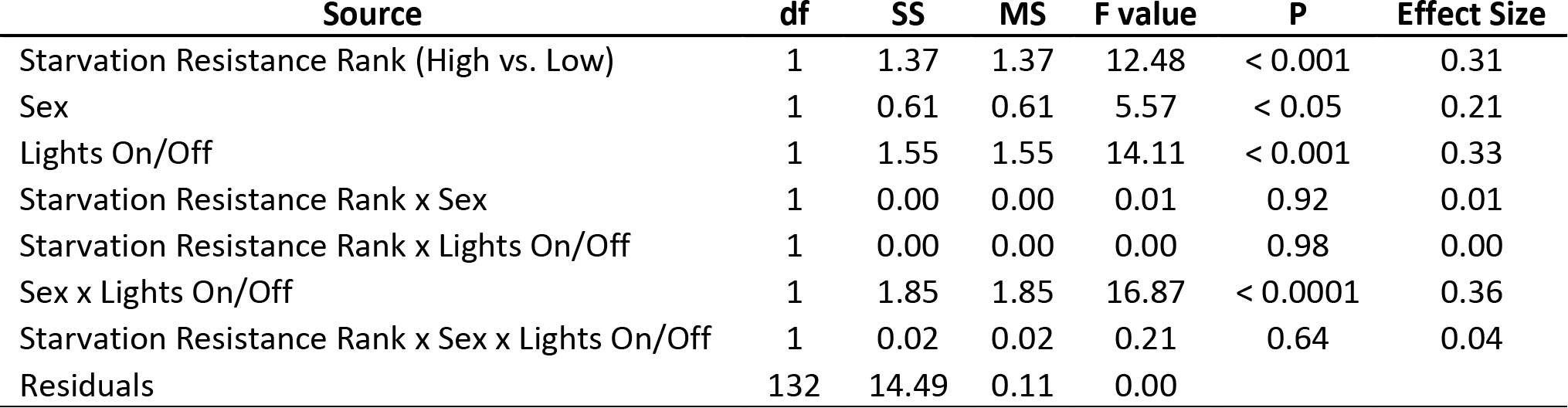
Analysis of variance of activity during the 24-hour period prior to the DAM (Drosophila Activity Monitor) starvation assay.

| Source | df | SS | MS | F value | P | Effect Size |
| --- | --- | --- | --- | --- | --- | --- |
| Starvation Resistance Rank (High vs. Low) | 1 | 1.37 | 1.37 | 12.48 | < 0.001 | 0.31 |
| Sex | 1 | 0.61 | 0.61 | 5.57 | < 0.05 | 0.21 |
| Lights On/Off | 1 | 1.55 | 1.55 | 14.11 | < 0.001 | 0.33 |
| Starvation Resistance Rank x Sex | 1 | 0.00 | 0.00 | 0.01 | 0.92 | 0.01 |
| Starvation Resistance Rank x Lights On/Off | 1 | 0.00 | 0.00 | 0.00 | 0.98 | 0.00 |
| Sex x Lights On/Off | 1 | 1.85 | 1.85 | 16.87 | < 0.0001 | 0.36 |
| Starvation Resistance Rank x Sex x Lights On/Off | 1 | 0.02 | 0.02 | 0.21 | 0.64 | 0.04 |
| Residuals | 132 | 14.49 | 0.11 | 0.00 |  |  |

**Table S8.**
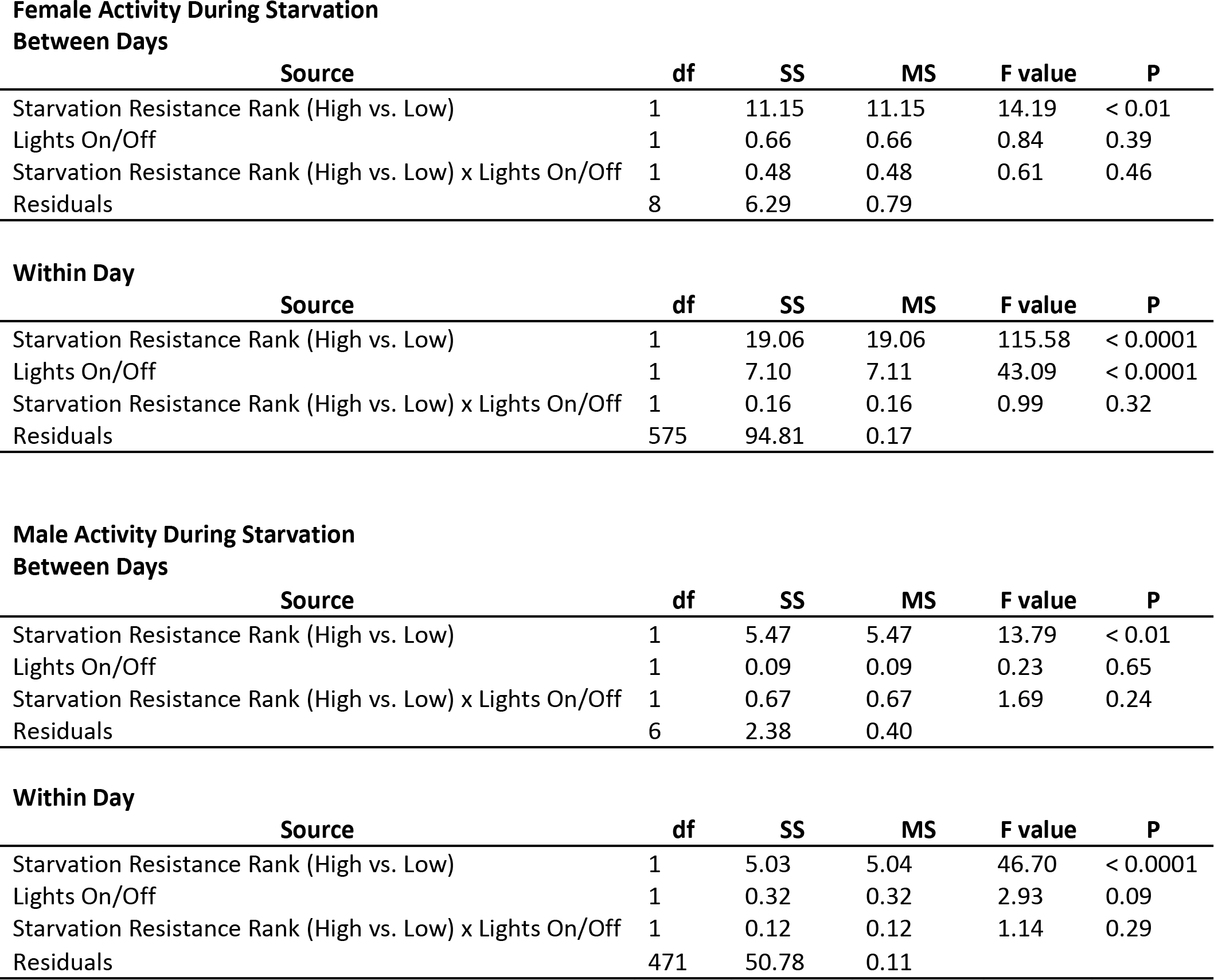
Repeated measures analysis of variance across days for activity during the DAM (Drosophila Activity Monitor) starvation assay for males and females.

**Table S9. Data from genes mapped to the region under QTL intervals for starvation resistance in the pA and pB DSPR mapping panels and triglyceride level in the pB DSPR mapping panel based on Flybase release version FB2018_1**. Highlighted genes indicate those previously identified in QTL mapping studies of starvation resistance.

**Table S10.**
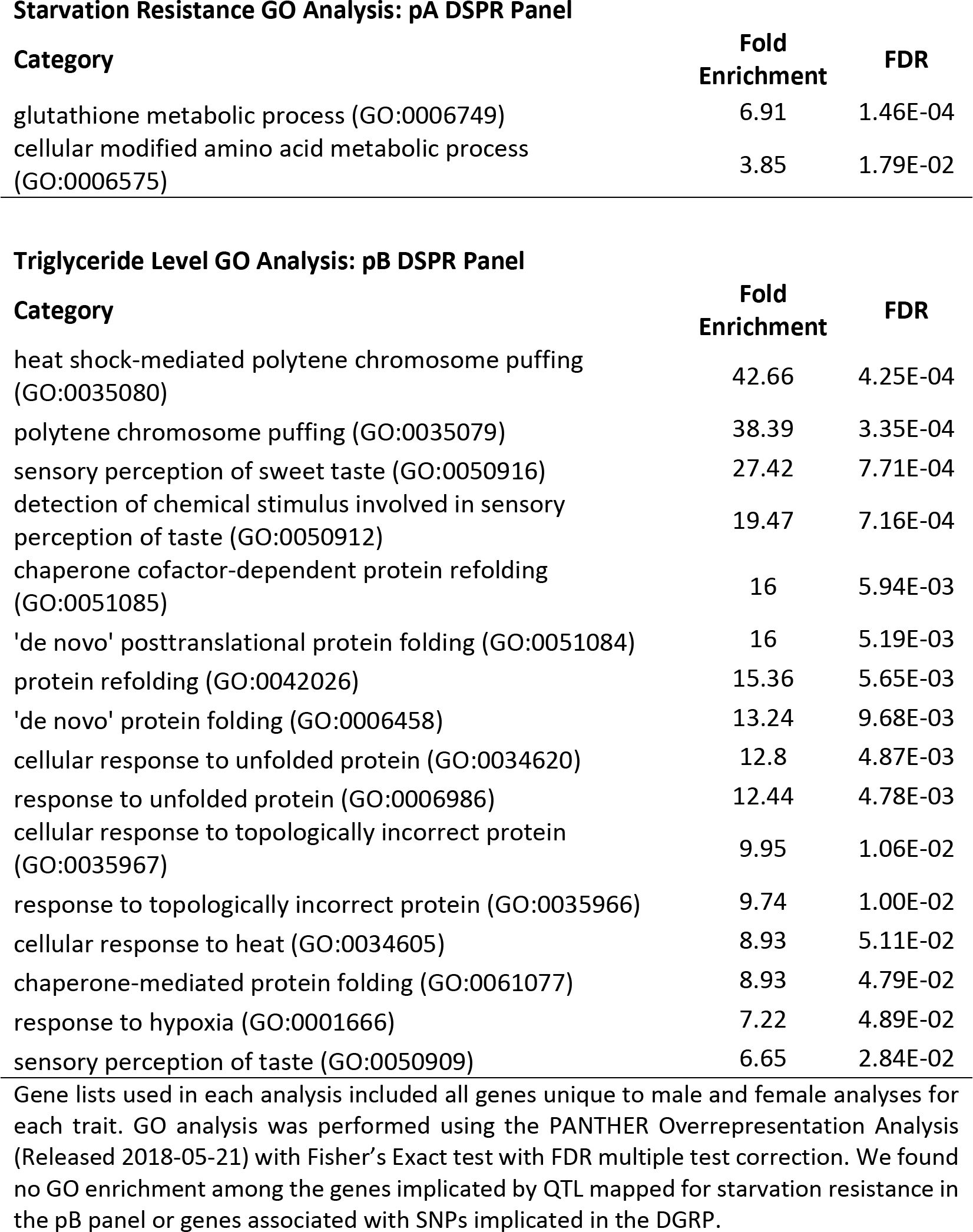
Gene ontology analysis of genes that are included within QTL intervals for starvation resistance and triglyceride level.

**Table S11. Data from GWA, generated from the DGRP Freeze 2.0 pipeline, based on Flybase release version FB2018_1.** All SNPs shown passed the P < 10^−5^ significance threshold; highlighted SNPs passed the FDR threshold of 0.2.

## Supplemental Text

**Text S1.** Starvation media recipe.

|  |  |
| --- | --- |
| 1000 ml | De-ionized water |
| 15 g | Agar |
| 12 ml | Propionic/Phosphoric acid mix* |
| 2 g | Tegosept (Genesee Scientific, Cat: 20-258) dissolved in 20 ml 95% EtOH |

\* Acid mix:

|  |  |
| --- | --- |
| 330 ml | De-ionized water |
| 259 ml | Propionic acid |
| 31 ml | o-Phosphoric acid (85%) |

Bring agar in water to a boil, reduce heat and simmer for 20 minutes. Remove from heat, stir in acid mix and tegosept. Cool slightly before pouring vials.

## Supplemental Files

File S1. All code associated with the bootstrapping analysis of SNPs associated with starvation resistance measured in the DGRP in this study, Mackay *et al.* (2012), and Everman and Morgan (2018).

**File S2**. *Description of each dataset associated with this study.*

\##### README for Datafiles (in RawData.zip) Accompanying Everman et al. 2019

\# DSPR_Data/:

### DSPR_1.txt

Tab-separated txt file of raw DSPR Starvation and Desiccation Resistance reported in hours for each fly per experimental vial.

Data are presented in Figure 1, Figure 3, Figure 6, Figure S2, Figure S5, Figure S7, Figure S11.

Column headers:

~~~
         Mapping.Panel = DSPR, Drosophila Synthetic Population Resource
         Trait = StarvationResistance or DesiccationResistance
         RIL.ID = Recombinant Inbred Line ID from the DSPR
         VialID = Unique number corresponding to each experimental vial
         Sex = (m) Male or (f) Female
         FlyID = Unique identifier for individual flies in each vial
         LifespanHrs = Lifespan in hours for each individual fly
~~~

### DSPR_2.txt

Tab-separated txt file of raw Founder Starvation Resistance reported in hours for each fly per experimental vial.

Data are presented in Figure 1.

Column headers:

~~~
         Trait = StarvationResistance
         DSPR.founder = DSPR founder line ID
         Sex = (m) Male or (f) Female
         RepVial = Vial replicate number
         FlyID = Unique identifier for individual flies in each vial
         LifespanHrs = Lifespan in hours for each individual fly
~~~

### DSPR_3.txt

Tab-separated txt file of raw female DSPR Triglyceride Levels reported per well. Data are presented in Figure 5, Figure 6, Figure S3.

Column headers:

~~~
         Mapping.Panel = DSPR, Drosophila Synthetic Population Resource
         Trait = TriglycerideLevel
         RIL.ID = Recombinant Inbred Line ID from the DSPR
         NumericPlateID = Unique number corresponding to each plate
         WellID = ID corresponding to each well of the plate
         SampleID = Unique identifier for samples
         TrueSerumTriglyConc = Triglyceride level per sample based on five females per sample
~~~

### DSPR_4.txt

Tab-separated txt file of raw DSPR Starvation Resistance reported in hours using Drosophila Activity Monitors (DAM).

Data are presented in Figure S7.

Column headers:

~~~
         Mapping.Panel = DSPR, Drosophila Synthetic Population Resource
         Trait = StarvationResistance
         RIL.ID = Recombinant Inbred Line ID from the DSPR
         StarvationClass = Categorical variable (HighStarvClass or LowStarvClass) based on data
         from DSPR_1.txt
         Sex = (M) Male or (F) Female
         MonitorID = Unique identifier for DAM
         MonitorTubeID = Unique identifier for each tube in each DAM
         LifespanHrs = Lifespan in hours for each fly
~~~

### DSPR_5.txt

Tab-separated txt file of DSPR Activity reported under non-stressful conditions using Drosophila Activity Monitors (DAM).

Data are presented in Figure 4.

Column headers:

~~~
         Mapping.Panel = DSPR, Drosophila Synthetic Population Resource
         RIL.ID = Recombinant Inbred Line ID from the DSPR
         Trait = Activity
         Sex = (m) Male or (f) Female
         StarvationClass = Categorical variable (HighStarvClass or LowStarvClass) based on data
         from DSPR_1.txt
         N = Number of flies tested
         ActLight.Mean = Mean activity levels under light conditions
         ActLight.SD = Standard deviation for activity levels under light conditions
         ActDark.Mean = Mean activity levels under dark conditions
         ActDark.SD = Standard deviation for activity levels under dark conditions
~~~

### DSPR_6.txt

Tab-separated txt file of DSPR Activity reported under starvation conditions using Drosophila Activity Monitors (DAM).

Data are presented in Figure S13.

Column headers:

~~~
         Mapping.Panel = DSPR, Drosophila Synthetic Population Resource
         RIL.ID = Recombinant Inbred Line ID from the DSPR
         LightStatus = (L) light or (D) dark
         SamplingPeriod = Day of experiment
         FemaleMeanActivity = Mean activity of females
         MaleMeanActivity = Mean activity of males
~~~

### DSPR_7.txt

Tab-separated txt file of DSPR Average Starvation Resistance reported in hours under different rearing density treatments.

Data are presented in Figure S1.

Column headers:

~~~
         Mapping.Panel = DSPR, Drosophila Synthetic Population Resource
         Trait = StarvationResistance
         RIL.ID = Recombinant Inbred Line ID from the DSPR
         Treatment = Density (60 eggs placed into rearing vials) or Population (females laid eggs
         for 1-2 days with visual assessment of density)
         TotalAdults = Total number of flies that emerged from each vial
         VialID = Unique identifier for each experimental vial
         N = Number of flies per experimental vial
         LifespanHrs = Average lifespan in hours for each experimental vial
~~~

### DSPR_8.txt

Tab-separated txt file of DSPR % survival on starvation media under different rearing density treatments.

Data are presented in Figure S1.

Column headers:

~~~
         Mapping.Panel = DSPR, Drosophila Synthetic Population Resource
         Trait = StarvationResistance
         RIL.ID = Recombinant Inbred Line ID from the DSPR
         Treatment = Density (60 eggs placed into rearing vials) or Population (females laid eggs
         for 1-2 days with visual assessment of density)
         TotalAdults = Total number of flies that emerged from each vial
         VialID = Unique identifier for each experimental vial
         ScreenpointHrs = Hour intervals at which flies in each vial were counted
         Survival% = Percent of flies in each vial that were alive at each screenpoint
~~~

### DSPR_9.txt

Tab-separated txt file of DSPR LOD score from QTL mapping analysis. Data are presented in Figure S14 and Figure S15.

Column headers:

~~~
         DSPR.Panel = pA or pB mapping panel from the DSPR
         Trait = StarvationResistance or TriglycerideLevels
         sex = (m) Male or (f) Female
         chr = Chromosome (X, 2L, 2R, 3L, 3R)
         Ppos = Position on chromosome based on assembly 5.0
         Gpos = Genetic position
         LOD = LOD score from QTL mapping analysis
~~~

### DSPR_10.txt

Tab-separated txt file of DSPR observed and estimated starvation resistance. Data are presented in Figure 7.

Column headers:

~~~
         DSPR.founder = pA or pB mapping panel from the DSPR
         ObservedStarvationResistance = Mean observed starvation resistance of each founder line
         EstimatedStarvationResistance = Weighted estimated mean starvation resistance of each founder line determined from QTL analysis
~~~

### DSPR_11.txt

Tab-separated txt file of DSPR starvation resistance and triglyceride level after accounting for variation due to haplotype at overlapping peaks.

Data are presented in Figure S16.

Column headers:

~~~
         FounderHaplotype = Predicted haplotype at overlapping QTL
         StarvationResistance = Mean starvation resistance
         TriglycerideLevel = Mean triglyceride level
         N_Starv = Number of RILs with the corresponding predicted founder haplotype for
         Starvation QTL
         N_Tri = Number of RILs with the corresponding predicted founder haplotype for
         Triglyceride QTL

##############################################################################
#################################################################
~~~

DGRP_Data/:

### DGRP_1.txt

Tab-separated txt file of raw DGRP Starvation Resistance reported in hours for each fly per experimental vial. Average response per sex and line was used in GWA.

Data are presented in Figure 2, Figure S4, Figure S5, Figure S8, Figure S12, used to generate Figure 17, Figure S18, Figure S19.

Column headers:

~~~
         Mapping.Panel = DGRP, Drosophila Genetic Reference Panel
         Trait = StarvationResistance
         RAL.ID = Line ID based on RAL identifier
         Bloomington.ID = Bloomington stock ID
         Sex = (M) Male or (F) Female
         VialID = Unique identifier for each experimental vial
         FlyID = Unique identifier for individual flies in each vial
         LifespanHrs = Lifespan in hours for each individual fly
~~~

### DGRP_2.txt

Tab-separated txt file of DGRP Average Starvation Resistance reported in hours in different environments and on different starvation media types.

Data are presented in Figure S9.

Column headers:

~~~
         Mapping.Panel = DGRP, Drosophila Genetic Reference Panel
         Trait = StarvationResistance
         RAL.ID = Line ID based on RAL identifier
         Environment = Flies were maintained at 25°C with a 12:12hr L:D cycle (25°C_12hr) or
         23°C with constant light (23°C_24hr)
         Media = Preservatives or NoPreservatives in the starvation media
         VialID = Unique identifier for each experimental vial
         N = Number of flies per experimental vial
         LifespanHrs = Average lifespan in hours for each experimental vial
~~~

### DGRP_3.txt

Tab-separated txt file of DGRP % survival in different environments and on different starvation media types.

Data are presented in Figure S9.

Column headers:

~~~
         Mapping.Panel = DGRP, Drosophila Genetic Reference Panel
         Trait = StarvationResistance
         RAL.ID = Line ID based on RAL identifier
         Media = Preservatives or NoPreservatives in the starvation media
         Environment = Flies were maintained at 25°C with a 12:12hr L:D cycle (25°C_12hr) or
         23°C with constant light (23°C_24hr)
         VialID = Unique identifier for each experimental vial
         ScreenpointHrs = Hour intervals at which flies in each vial were counted
         Survival% = Percent of flies in each vial that were alive at each screenpoint
~~~

### DGRP_4.txt

Tab-separated txt file of DGRP Starvation Resistance for lines shared between this study, Mackay et al. 2012, and Everman and Morgan 2018.

Data are presented in Figure S8, Figure S10.

Column headers:

~~~
         RAL.ID = Line ID based on RAL identifier
         Sex = (M) Male or (F) Female
         LifespanHrs_EvermanetAl.2019 = Mean starvation resistance reported in this study
         LifespanHrs_MackayEtAl.2012 = Mean starvation resistance reported in Mackay et al. 2012
         LifespanHrs_Everman&Morgan.2018 = Mean starvation resistance reported in Everman and Morgan 2018
~~~

### DGRP_5.txt

Tab-separated txt file of bootstrap results of the sign of SNPs across DGRP data collected in this study, Mackay et al. 2012 and Everman and Morgan 2018.

These data are compiled from original bootstrap files; code presented in File S1 is formatted to read each file by DataID.

Data are presented in Figure 8.

Column headers:

~~~
         %SNPs_SameSign = Percent of SNPs that have the same sign between studies following boostrap analysis of random samples of SNPs.
         Density = Density calculated from original bootstrap files
         Sex = Female or Male
         DataID = Dataset ID for plotting in File S1
         Comparison = Direction of comparison of SNPs (ThisStudy_vs_MackayEtAl2012,
ThisStudy_vs_Everman&Morgan2018, Everman&Morgan2018_vs_ThisStudy,
Everman&Morgan2018_vs_MackayEtAl2012, MackayEtAl2012_vs_ThisStudy, or
MackayEtAl2012_vs_Everman&Morgan2018)
~~~

### DGRP_6.txt

Tab-separated txt file of adjusted mean starvation resistance DGRP data collected in this study, Mackay et al. 2012 and Everman and Morgan 2018.

These data are compiled from original GWA-generated files; code presented in File S1 is formatted to read each file by Study.

Data are used in bootstrap analysis.

Column headers:

~~~
         RAL.ID = Line ID based on RAL identifier
         AdjustedMeanStarvationResistance = Adjusted mean phenotype from GWA of each study
         Study = Study in which the original starvation resistance data was collected (ThisStudy,
         MackayEtAl2012, or Everman&Morgan2018)
         Sex = (f) female or (m) male
~~~

### DGRP_7.txt

Comma-separated txt file of SNP frequencies for bootstrap analysis, used in File S1, generated from dgrp.t.txt in code file.

Data are used in bootstrap analysis.

Column headers:

~~~
         rAF = Reference allele frequency
         aAf = Alternate allele frequqncy
         SNP = SNP ID
~~~

### DGRP_8.txt

Large space-separated matrix of SNP calls for bootstrap analysis, used in File S1, called as dgrp.t.txt in code file. Data originally available from dgrp2.gnets.ncsu.edu.

Data are used in bootstrap analysis.

Column headers:

~~~
         id = Line ID based on RAL identifier, formatted “line_XXX”
         *Remaining Columns: SNP id
~~~

Additional Files:

/GWAS_AVG_Starvation/

Contains output files for the average starvation resistance calculated for overlapping

DGRP lines between this study, Mackay et al. 2012 and Everman and Morgan 2018.

/GWAS_SJM_Starvation/

Contains output files for the average starvation resistance from this study.

